# High-density transposon mutagenesis in *Mycobacterium abscessus* identifies an essential penicillin-binding lipo-protein (PBP-lipo) involved in septal peptidoglycan synthesis and antibiotic sensitivity

**DOI:** 10.1101/2021.07.01.450732

**Authors:** Chidiebere Akusobi, Bouchra S. Benghomari, Junhao Zhu, Ian D. Wolf, Shreya Singhvi, Charles L. Dulberger, Thomas R. Ioerger, Eric J. Rubin

## Abstract

*Mycobacterium abscessus* (*Mab*) is a rapidly growing non-tuberculous mycobacterium (NTM) that causes a wide range of infections. Treatment of *Mab* infections is difficult because the bacterium is intrinsically resistant to many classes of antibiotics. Developing new and effective treatments against *Mab* requires a better understanding of the unique vulnerabilities that can be targeted for future drug development. To achieve this, we identified essential genes in *Mab* by conducting transposon-sequencing (TnSeq) on the reference *Mab* strain ATCC 19977. We generated ∼51,000 unique transposon mutants and used this high-density library to identify 362 essential genes for *in vitro* growth. To investigate species-specific vulnerabilities in *Mab*, we further characterized *MAB_3167c*, a predicted penicillin-binding-lipoprotein (PBP-lipo) that is essential in *Mab* and non-essential in *Mycobacterium tuberculosis* (*Mtb*). We found that PBP-lipo primarily localizes to the subpolar region and later to the septum as cells prepare to divide. Depletion of *Mab* PBP-lipo causes cells to elongate, develop ectopic branches, and form multiple septa. Knockdown of PBP-lipo along with PbpB, DacB1, and a carboxypeptidase, MAB_0519 lead to synergistic growth arrest. In contrast, these genetic interactions were absent in the *Mtb* model organism, *Mycobacterium smegmatis*, indicating that the PBP-lipo homologs in the two species exist in distinct genetic networks. Finally, repressing PBP-lipo sensitized the reference strain and 11 *Mab* clinical isolates to several classes of antibiotics, including the β-lactams, ampicillin and amoxicillin by greater than 128-fold. Altogether, this study presents PBP-lipo as a key enzyme to study *Mab* specific processes in cell wall synthesis and importantly positions PBP-lipo as an attractive drug target to treat *Mab* infections.

## Introduction

*Mycobacterium abscessus* (*Mab*) is the most common cause of human disease among the rapidly growing non-tuberculous mycobacteria (NTM). It causes a wide range of illnesses including lung, skin and soft-tissue infections, as well as disseminated disease (1–3). While the incidence of *Mab* infections is rising worldwide, treating *Mab* infections remains difficult. The bacterium has intrinsic and acquired resistance mechanisms to many classes of antibiotics, including the standard drugs used to treat tuberculosis (4, 5). The current treatment regimen for *Mab* requires taking a combination of multiple antibiotics for up to 18 months and is often associated with severe toxicity and routinely ends in treatment failure (6). Thus, there is an urgent need for new and effective drugs to treat the emerging global public health threat of *Mab* infections (7).

Developing novel antibiotic treatments for *Mab* requires a better understanding of essential biological processes in the bacterium that can be targeted. Identifying essential genes is the first step in this process, and by combining transposon-mutagenesis with massive parallel sequencing, transposon-sequencing (TnSeq) has been an effective tool to determine gene essentiality on a genome-wide scale in other microbes (8). However, while many TnSeq screens have been conducted in *Mycobacterium tuberculosis* (*Mtb*) (9), up until recently, only two *Mab* TnSeq screens have been published (10, 11). While both *Mab* TnSeq screens provided tools and important insights into *Mab* biology, neither utilized a mutant library with high enough density to comprehensively identify essential genes. This changed with the recent publication from Rifat *et al*, who generated robust high-density transposon libraries of *Mab* and identified a comprehensive list of genetic elements essential for *in vitro* growth (12). Identification of these essential genes serves as an excellent starting point to classify attractive targets for drug discovery.

One major target of antibiotics is the mycobacterial cell wall whose biogenesis is essential for bacterial growth (13, 14). The structure of the mycobacterial cell wall is unique among bacteria because it is composed of three macromolecules: peptidoglycan (PG), arabinogalactan, and mycolic acids. Together, these layers envelop mycobacteria with a thick, waxy coat that forms a permeability barrier to many drugs (15, 16). The foundational layer is comprised of peptidoglycan (PG), which consists of glycan sugars cross-linked by short peptides forming a continuous, net-like structure that maintains cell shape and prevents rupture. Mycobacteria build new PG at the cell poles aided by Class A penicillin binding proteins (PBPs) that perform two reactions: a transglycosylation reaction that polymerizes PG monomers to existing glycan strands and a transpeptidation reaction that cross-links glycan strands to each other (17). Class B PBPs solely perform the cross-linking reactions (18, 19). A hallmark of PBPs is that they are inhibited by β-lactam antibiotics; however, these antibiotics are typically not used to treat *Mab* infections due to high levels of resistance and the presence of β-lactamases in the genome (20).

To identify essential genes for *in vitro* growth and classify potential new drug targets in *Mab*, we performed TnSeq. We generated high-density transposon libraries of ∼51,000 mutants in the reference strain, *M. abscessus subsp. abscessus* ATCC 19977 and identified 362 genes required for *in vitro* growth. We next identified essential genes in *Mab* that were non-essential in *Mtb* in order to study species-specific vulnerabilities in *Mab*. From this list of genes, we further characterized *MAB_3167c*, which encodes PBP-lipo, a conserved Class B PBP. We found that PBP-lipo is required for normal peptidoglycan (PG) synthesis and cell division. Knockdown of PBP-lipo sensitizes *Mab* to commonly-used β-lactams and other classes of antibiotics. None of these phenotypes were present in the *Mycobacterium smegmatis (Msm)* and *Mtb* PBP-lipo knockouts. Thus, we find that PBP-lipo has a unique essential function in *Mab* and is a potential drug target for treating *Mab* infections.

## Results

### Identification of *M. abscessus* ATCC 19977 essential genes for *in vitro* growth

To identify essential genes in *Mab*, we performed TnSeq on *M. abscessus subsp. abscessus* ATCC 19977 (hereafter referred to as *Mab*) using the ϕMycoMar T7 phage carrying the Himar1 transposon for transduction. This transposon inserts into ‘TA’ dinucleotide sites and disrupts gene function (Figure 1A) (21). We transduced three independently grown cultures of *Mab*, yielding triplicate libraries with saturations of 54.2%, 62.3%, and 64.1% for Library 1, 2, and 3 respectively (Supplemental Table 1). The average Pearson r^2^ correlation of mean read counts for each gene across 2 libraries was 0.85 indicating good correlation among the libraries (Supplemental Figure 1). The transposon insertions across the three libraries were mapped onto the *Mab* genome (Figure 1B). Across three libraries, we generated approximately 51,000 unique transposon mutants, as defined by sites of unique transposon insertions across all three libraries. Of note, the *Mab* libraries shared the same non-permissive ‘TA’ sites for transposon insertion as has been described in other mycobacterial species (22).

**Figure 1.**
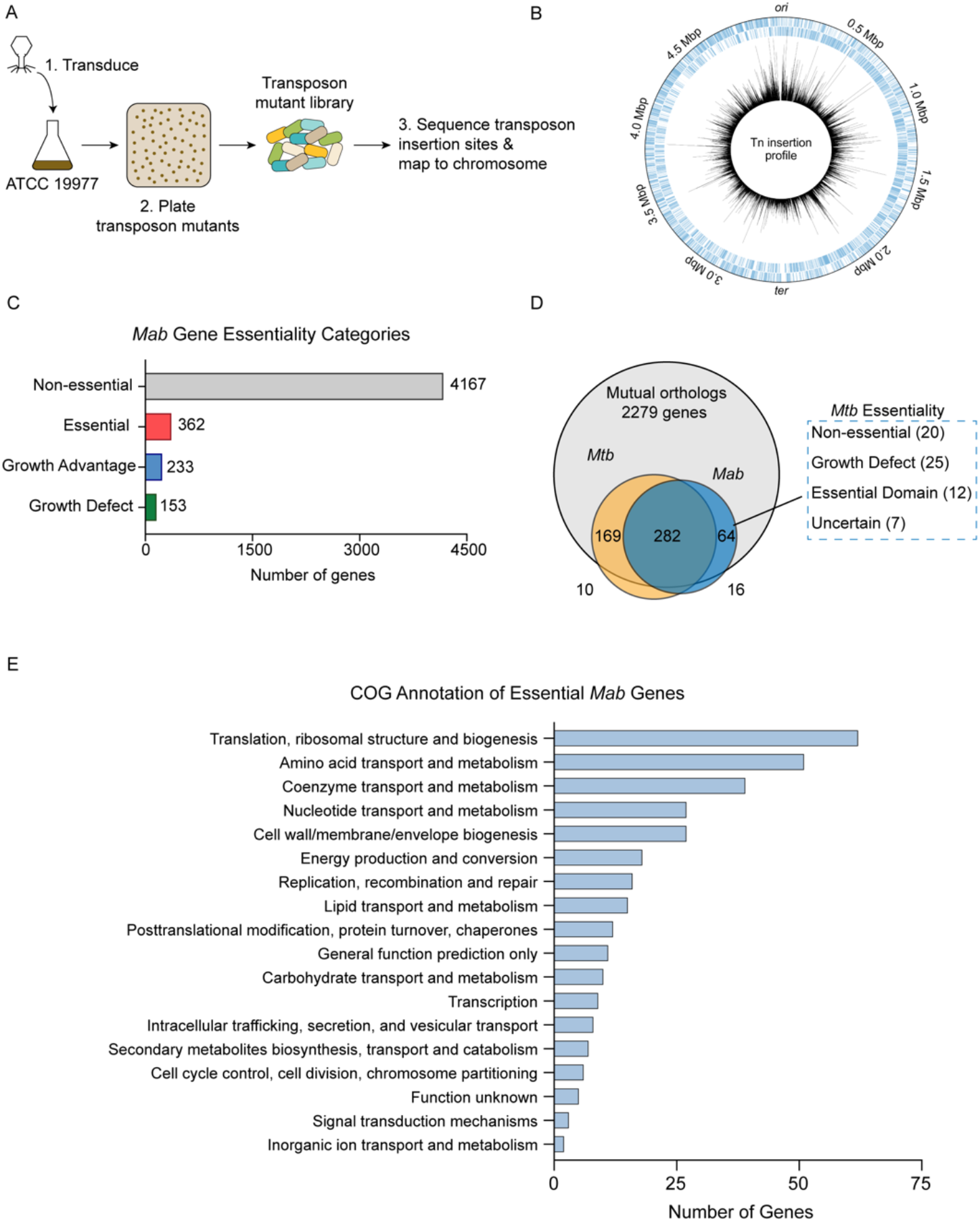
Overview of TnSeq Analysis. **(A)** Schematic of TnSeq protocol. Phage carrying Himar1 transposon was used to transduce *Mab subsp. abscessus* ATCC 19977 cultures. Over 51,000 independent transposon mutants were generated across three libraries **(B)** Location of transposon insertions in *Mab* genome. Black lines represent the average number of transposon insertions per gene across the three replicates. **(C)** Breakdown of gene essentiality categories as determined by the Hidden Markov Model (HMM) in TRANSIT **(D)** Essentiality comparison of mutual orthologs between *Mab* and *Mtb*. **(E)** Clusters of Orthologous Group (COG) categories of essential genes in *Mab*.

The high density of these libraries allowed us to identify essential *Mab* genes using the Hidden Markov Model (HMM) algorithm implemented in TRANSIT (23). HMM identified 362 genes as essential, 4167 as non-essential, 153 as producing a growth defect when disrupted, and 233 as causing a growth advantage when disrupted (Figure 1C, Supplemental Table 2). As expected, many of the genes identified as essential were involved in key biological processes including protein translation, amino acid metabolism, and cell wall biogenesis based on Clusters of Orthologous Groups (COG) analysis (Figure 1E).

### Identification and validation of uniquely essential genes in *Mab*

Understanding differences in essential genes between *Mab* and *Mtb* might uncover new ways of specifically targeting *Mab*. Thus, we identified *Mab-*specific essential genes by comparing gene essentiality between mutual orthologs of *Mab* and *Mtb.* We used *Mtb* essentiality data generated from a comprehensive analysis using 14 high-quality TnSeq datasets (9). Of the 2279 mutual orthologs between *Mab* and *Mtb*, 282 genes were essential in both species. We focused on the mutual orthologs that that were essential in *Mab,* but non-essential in *Mtb*. These genes could potentially represent *Mab* specific drug targets. Exactly 20 genes fit into this category, which we refer to as “uniquely essential” (Figure 1D) (Supplemental Table 3).

To understand the quality of predictions by TnSeq, we validated the essentiality of a subset of genes using an anhydrotetracycline (ATc) inducible CRISPRi system developed for use in *Mtb*(24). Expression of a non-targeting sgRNA control in this system had no impact on cell proliferation (Figure 2A). In contrast, repressing canonical essential genes *rpoB, gyrB,* and *secY* impaired cell growth in both liquid broth (Figure 2A) and solid media (Supplemental Figure 2A), indicating that the CRISPRi system developed in *Mtb* functions in *Mab*.

**Figure 2.**
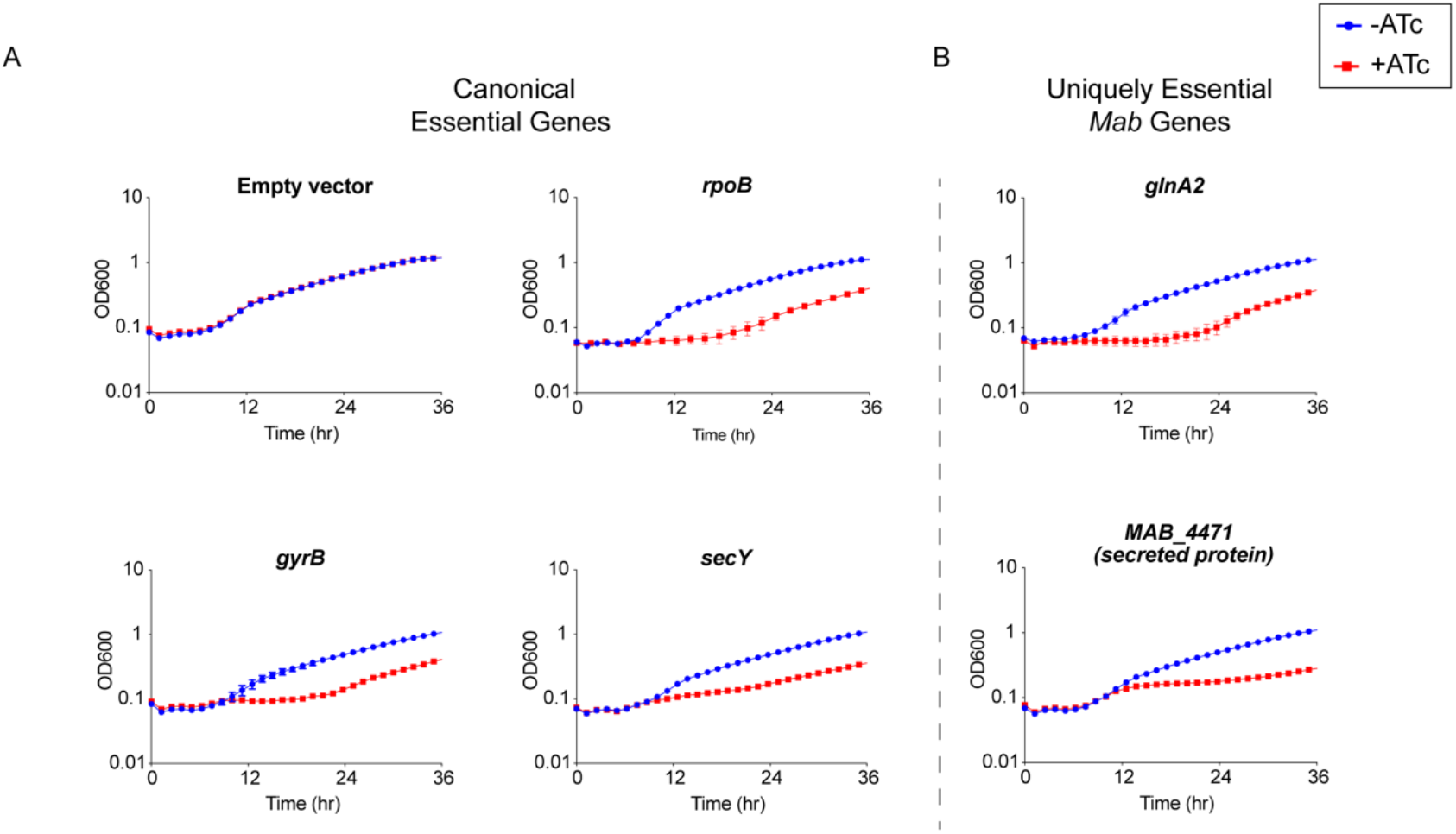
Validation of Essential Genes. **(A)** Growth curve of *Mab* strains with sgRNAs targeting canonical essential genes. **(B)** Growth curve of *Mab* strains with sgRNAs targeting uniquely essential *Mab* genes.

We next validated the essentiality of a subset of the 20 uniquely essential genes in *Mab*: *glnA2, icd2, sdhA,* and *MAB_4471*. CRISPRi knockdown of these uniquely essential genes impaired growth in liquid media (Figure 2B) as well as solid media (Supplemental Figure 2B). These results confirm that at least a subset of uniquely essential *Mab* genes identified by TnSeq are required for optimal *in vitro* growth. The slight rise in OD_600_ later in the time course is consistent with the emergence of escape mutants in the CRISPRi system, which has been previously described (25, 26).

### PBP-lipo is uniquely essential in *Mab*

One of the uniquely essential genes we identified was *MAB_3167c*, which encodes a hypothesized Class B penicillin-binding lipoprotein (PBP-lipo). PBP-lipo is predicted to have a single N-terminal transmembrane helix and a transpeptidase domain. We found that *MAB_3167c* lacked transposon insertions across all three replicate libraries (Figure 3A). This finding corroborates data from previously generated *Mab* transposon libraries that show lack of insertions in *MAB_3167c* (10, 12). Comparing the essentiality of PBP enzymes in *Mab* with their homologs in *Mtb* revealed that *MAB_3167c* is unique in being the sole PBP whose essentiality differs between the two species. Specifically, PBP-lipo is essential in *Mab* and non-essential in *Mtb* (Figure 3B).

**Figure 3.**
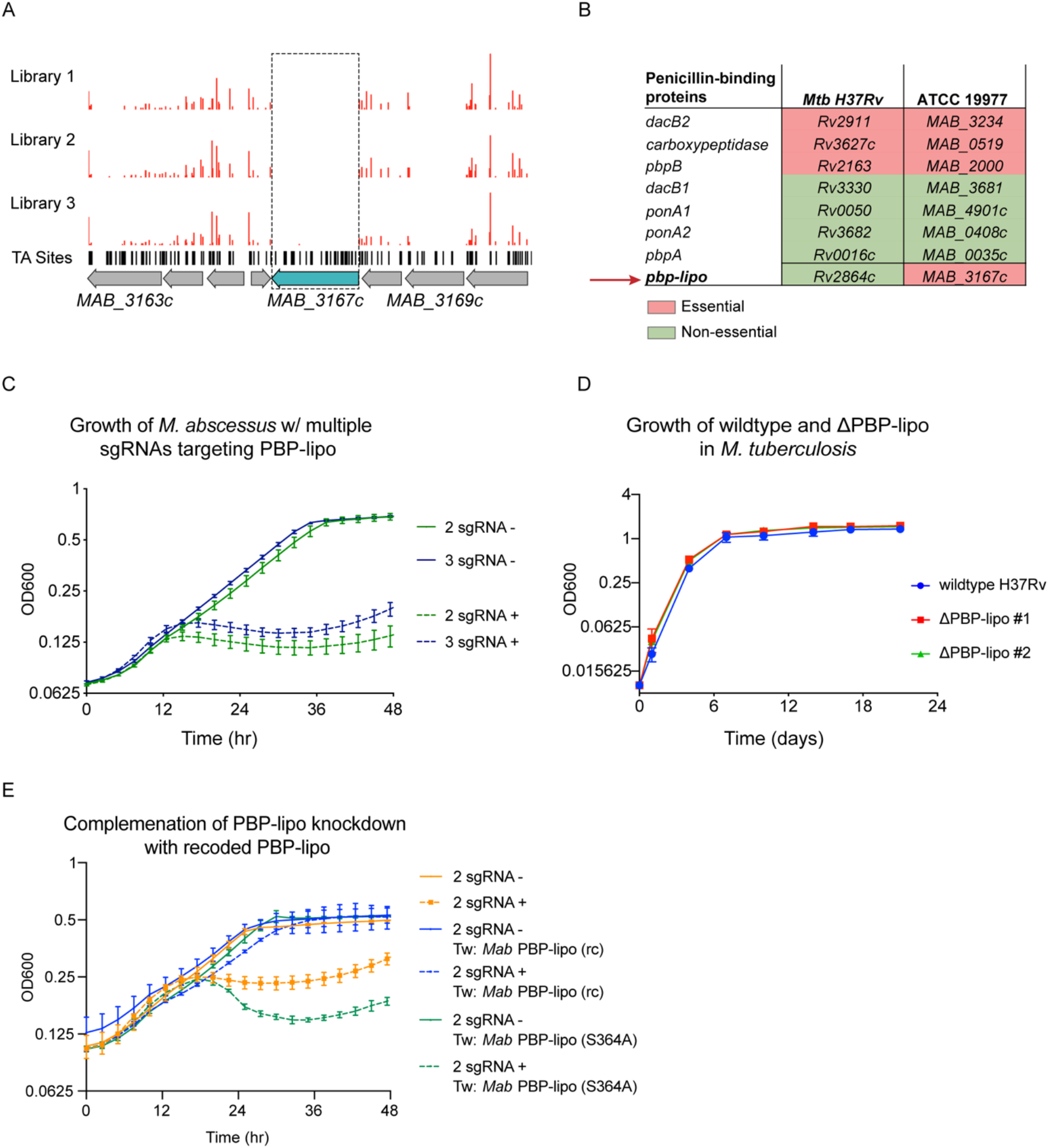
PBP-lipo is uniquely essential in *Mab* and non-essential in *Mtb*. **(A)** Transposon-insertion profile of PBP-lipo across three replicate transposon mutant libraries. Red bars indicate the number of transposon insertions. Horizontal black lines indicate ‘TA’ sites, where the transposon inserts. Genes are schematized with gray arrows, with PBP-lipo colored in blue. Dotted box demarcates lack of insertions in PBP-lipo. **(B)** Comparison of essentiality for PBP orthologs between *Mtb* and *Msm*. **(C)** Growth of *Mab* cultures transformed with CRISPRi plasmid carrying either 2 or 3 sgRNAs targeting PBP-lipo. “-” symbol indicates cultures were grown without ATc. “+” symbol indicates cultures were grown with ATc. **(D)** Growth of wildtype *Mtb* and PBP-lipo knockout strains. **(E)** Growth of *Mab* with native PBP-lipo knockdown complemented by recoded (rc) PBP-lipo, which sgRNAs can no longer bind. PBP-lipo (rc) (S364A) is a recoded and catalytically inactive version of the enzyme.

To validate that PBP-lipo is required for *in vitro* growth, we used the CRISPRi system to induce knockdown using constructs encoding 1, 2, or 3 sgRNAs targeting *MAB_3167c* (Supplemental Figure 3A). The single, double, and triple sgRNA constructs allowed for the fine tuning of the magnitude of PBP-lipo knockdown, as measured by RT-qPCR (Supplemental Figure 3B). We noted that the empty vector control (0 sgRNA) and single sgRNA construct had no detectable effect on the growth of *Mab* (Supplemental Figure 3C). In contrast, when *MAB_3167c* was targeted with either 2 or 3 sgRNAs, the growth of *Mab* was inhibited, indicating that PBP-lipo is indeed required for optimal *in vitro* growth (Figure 3C). We also observed inhibition of cell growth on solid media where knockdown of PBP-lipo by 2 sgRNA and 3 sgRNA vectors led to a 1000-fold reduction in colony forming units (CFU) (Supplemental Figure 3D). In contrast to *Mab*, when we deleted the gene encoding PBP-lipo in *Mtb*, *Rv2864c*, the growth of the knockout and wild-type strains was indistinguishable (Figure 3D). Similarly, the PBP-lipo homolog in *Msm, MSMEG_2584c*, when knocked out grew identically to wildtype with no growth defect (Supplemental Figure 4).

To complement the PBP-lipo knockdown growth defect phenotype in *Mab*, we recoded a version of PBP-lipo with the protospacer adjacent motif (PAM) and sgRNA binding sequences mutated while preserving the amino acid sequence. This prevents sgRNAs from binding the exogenous PBP-lipo copy present on the integrated plasmid. When constitutively expressed, this recoded PBP-lipo rescued growth of the native PBP-lipo knockdown (Figure 3E). In contrast, a recoded allele that was predicted to be catalytically inactive (S364A) did not rescue growth. All together, this data demonstrates that PBP-lipo’s predicted transpeptidase activity is required for *Mab* optimal growth, while PBP-lipo’s activity in *Mtb* and *Msm* is non-essential.

### PBP-lipo is required to maintain normal cell elongation and division

A hallmark of PBPs is that they bind β-lactam antibiotics. Therefore, to test whether PBP-lipo is an active PBP in *Mab*, we determined whether PBP-lipo binds bocillin FL, a fluorescent analog of penicillin (27). Using a strain that constitutively expresses a strep-tagged PBP-lipo, we identified a fluorescent bocillin FL band at the same molecular weight as PBP-lipo, which was confirmed to be PBP-lipo by Western blot (Supplemental Figure 5A). These data suggest that PBP-lipo binds bocillin FL *in vivo* and is an active PBP. Interestingly, in wildtype *Mab,* we did not detect an obvious PBP-lipo band at the predicted molecular weight of PBP-lipo, 63 kDa, suggesting that native expression levels of PBP-lipo is either low or PBP-lipo is expressed at distinct periods during the cell cycle.

We also tested whether PBP-lipo was indeed a lipoprotein. Based on its amino acid sequence, PBP-lipo has a N-terminal signal sequence that targets the protein for localization to the periplasm. This is followed by a conserved cysteine residue, which is the canonical site of lipid modification (28). To determine if PBP-lipo is lipidated, we performed a lipoprotein extraction protocol to isolate lipoproteins in *Mab* (29). By Western Blot, we successfully detected the presence of PBP-lipo in the lipoprotein fraction (Supplemental Figure 5B). PBP-lipo being present also in the non-lipoprotein fraction is not surprising as similar results were observed in early lipoprotein extraction experiments performed in *Mtb* (30)

In mycobacteria, the PG layer is critical for maintaining normal cell shape and altering PBP expression often results in abnormal cell morphology. Thus, we studied the morphology of *Mab* when PBP-lipo was repressed and discovered dramatic morphological changes. *Mab* cells elongate, branch inappropriately, and form ectopic poles of active growth, as evidenced by staining with NADA (3-[(7-Nitro-2,1,3-benzoxadiazol-4-yl)amino]-D-alanine), a fluorescent-D-amino acid (FDAA) that binds areas of active PG synthesis (31) (Figure 4A). PBP-lipo knockdown significantly increased the number of cells with ectopic branches (Supplemental Figure 6B). Furthermore, during normal growth of *Mab*, a small percentage of cells contain one mid-cell FDAA band, indicating active PG synthesis at the site of septation. However, when PBP-lipo is knocked down, we occasionally find cells with 2 or 3 bands mid-cell, as exemplified in the 2 sgRNA + panel (Figure 4A). PBP-lipo knockdown strains were all significantly longer than their uninduced controls, with the longest average cell length belonging to the strain with the strongest PBP-lipo knockdown (Figure 4B). Of note, while the 0 sgRNA +/- cells are morphologically indistinguishable, there is a statistically significant difference in the mean cell lengths, likely due to the effect of ATc on cells (Figure 4B). Interestingly, cell width did not change among the strains (Supplemental Figure 6A). In comparison, in *Msm* and *Mtb*, there were no significant differences in morphology and cell length between the wildtype and PBP-lipo knockout strains (Supplemental Figure 7). The dramatic changes in *Mab* morphology when PBP-lipo is repressed suggests proper expression levels of this enzyme is required for maintaining normal elongation and division in *Mab*.

**Figure 4.**
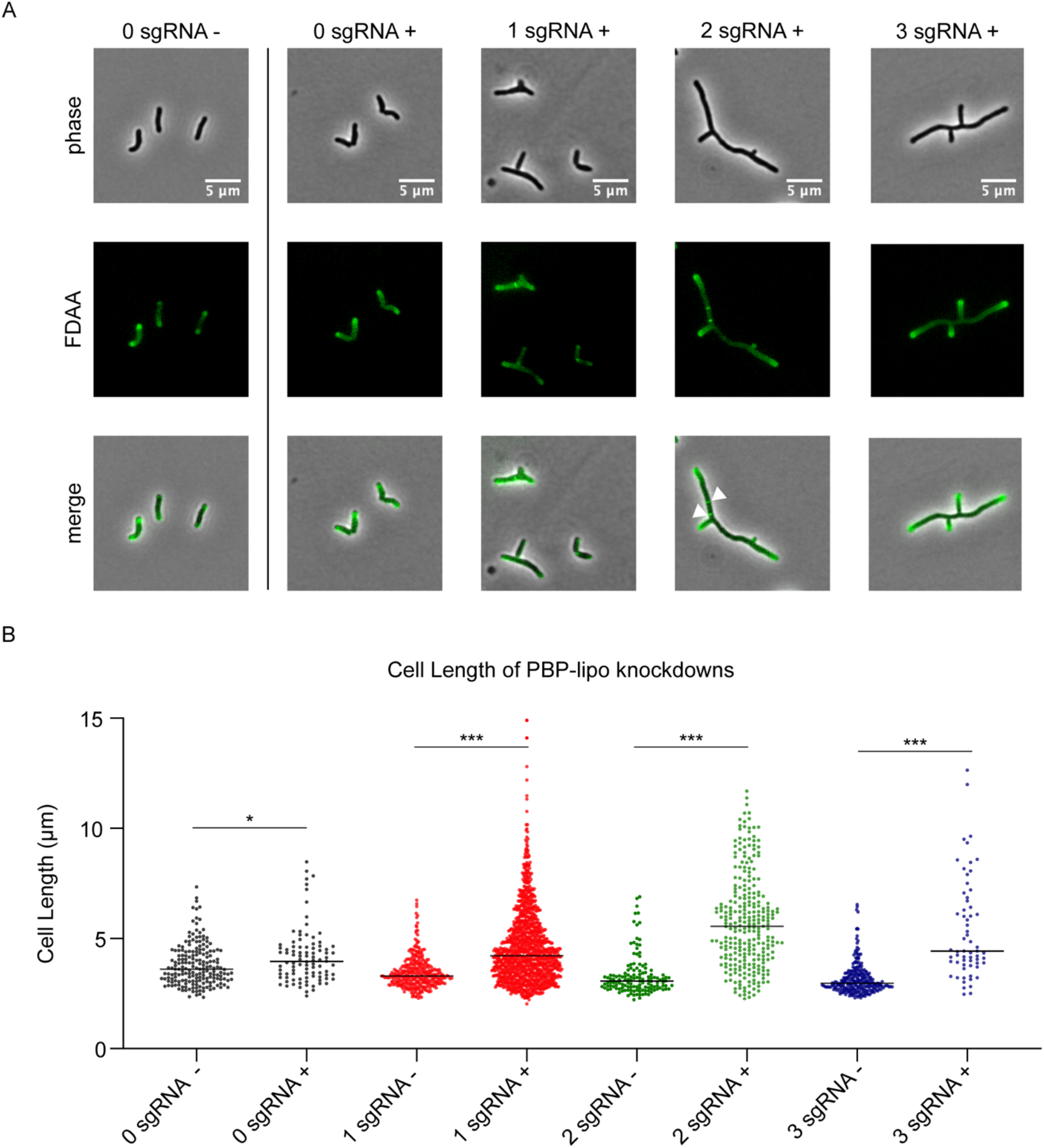
Knockdown of PBP-lipo disrupts cell morphology. **(A)** Microscopy images of PBP-lipo knockdown cultures. *Mab* strains carrying CRISPRi plasmids with either 0, 1, 2, or 3 sgRNAs targeting PBP-lipo. Arrows indicate sites of multiple septa formation. **(B)** Cell lengths of uninduced and induced strains. Measurements were obtained by GEMATRIA and MOMIA image analysis pipelines (35). Student’s *t test* used to calculate the statistical difference in mean cell lengths. *** = p-val < 0.0001. * = p-val < 0.05.

### PBP-lipo localizes to the septum after FtsZ

The morphological defects caused by PBP-lipo knockdown were reminiscent of septal factor depletions observed in *Msm* (32). Thus, we hypothesized that PBP-lipo localizes to the septum. To test this hypothesis, we N-terminally tagged PBP-lipo with mRFP, which has been previously shown to function in the mycobacterial periplasm (33). We cloned mRFP after the predicted N-terminal signal sequence of PBP-lipo with linkers on the 5’ and 3’ ends of the fluorescent protein (Supplementary Figure 8A). We next validated that this construct produced a full-length fusion protein using Western Blot, and detected the correctly sized 91kDa fusion product (Supplementary Figure 8B). Finally, we tested the functionality of the mRFP-PBP-lipo fusion protein by using recoded and non-recoded versions of the construct. The recoded version had a single sgRNA binding site on PBP-lipo mutated (Supplementary Figure 8A). As expected, knockdown of native PBP-lipo lead to growth inhibition while expression of the non-recoded fusion protein partially complemented growth. Reassuringly, the recoded mRFP-PBP-lipo fully complemented the growth defect from the native PBP-lipo knockdown indicating that the fusion protein is indeed functional.

Given the morphological defects in the PBP-lipo knockdown, we hypothesized that PBP-lipo localized to the septum. To test this, we imaged PBP-lipo simultaneously with FtsZ, which forms the Z-ring at the divisome. FtsZ is the first protein to localize to the septum and recruits other members of the divisome complex to aid in the formation of the septum and eventual division of the cell (34). To visualize FtsZ, we C-terminally tagged it with mNeonGreen (mNG) and expressed the fusion protein using a weak promoter as higher levels of expression are toxic. After introducing both constructs into wildtype *Mab*, we visualized the fusion proteins, which were expressed in the presence of wildtype PBP-lipo. Using fluorescence microscopy, we discovered that both FtsZ and PBP-lipo localized to the mid-cell, likely in the growing septum. We also identified cells with septal co-localization of both PBP-lipo and FtsZ (Figure 5A).

**Figure 5.**
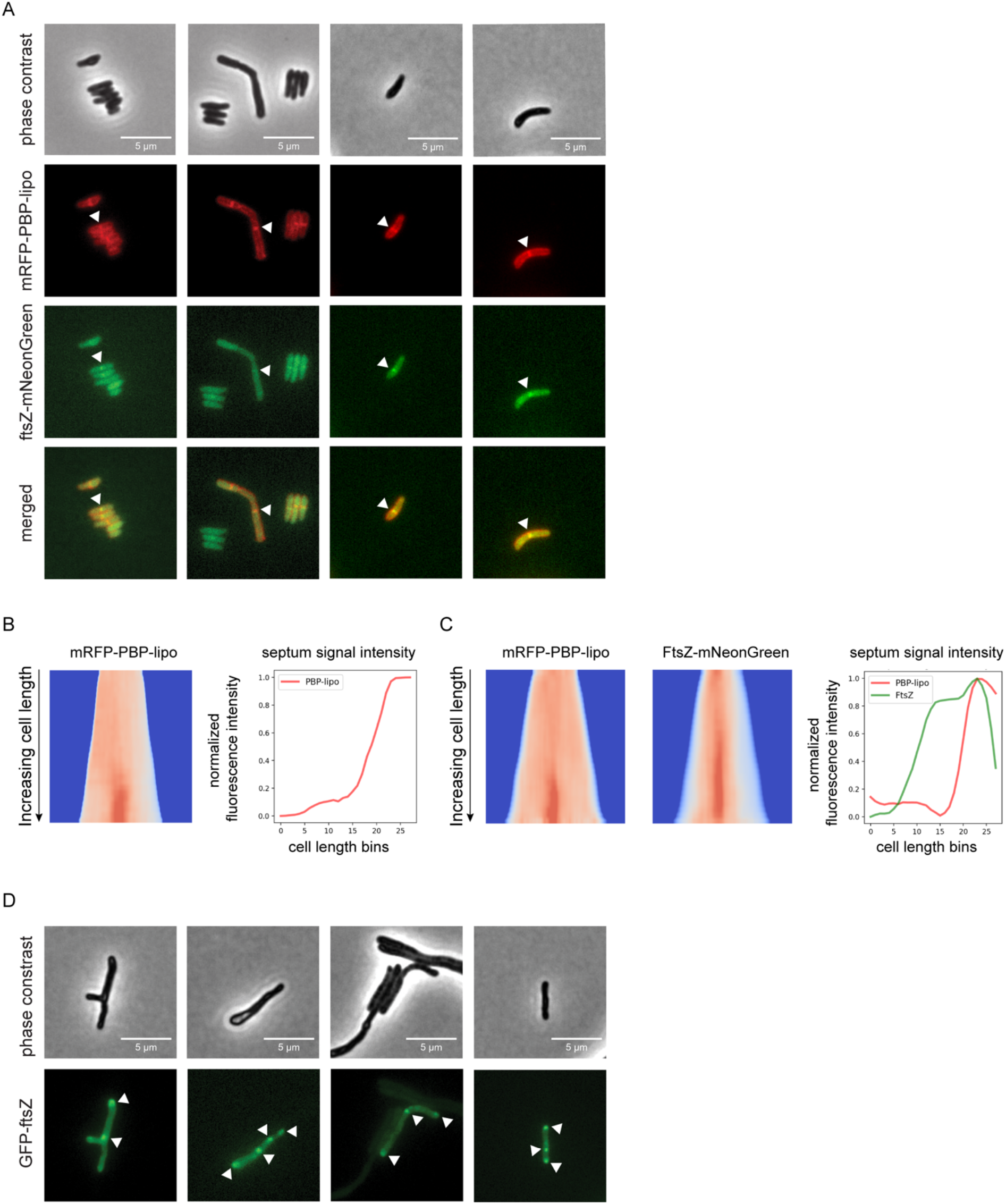
Knockdown of PBP-lipo disrupts formation of FtsZ rings. **(A)** Microscopy images of N-terminally tagged mRFP-PBP-lipo and C-terminally tagged FtsZ-mNeonGreen. ‘Merged’ images show overlay of red and green channels **(B)** (left) Demograph of mRFP-PBP-lipo. (right) Quantification of septal fluorescence signal across increasing cell lengths **(C)** (left) Demograph of mRFP-PBP-lipo and FtsZ-mNeonGreen. (right) Fluorescence signal arranged by increasing cell length **(D)** Images of GFP-FtsZ expressed from its natural promoter in the setting of PBP-lipo knockdown.

The resulting images were analyzed using the recently published MOMIA and GEMATRIA programs, which quantify the fluorescent signal across cells normalized by cell length (35). We first analyzed cells that expressed solely mRFP-PBP-lipo and demonstrated that mid-cell localization of PBP-lipo was only present in the longest cells in the population (Figure 5B). These data demonstrate that as cells elongate, PBP-lipo localizes to the septum and is therefore likely involved in septum formation and cell division. We next applied this analysis to cells that co-expressed mRFP-PBP-lipo and FtsZ-mNG. We found that FtsZ localizes to the septum earlier in the cell life cycle as demonstrated by the higher septal fluorescence signal intensity in shorter cells (Figure 5C). In contrast, PBP-lipo localizes to the septum when cells are longer and after FtsZ has already localized to the septum (Figure 5C). This sequential recruitment of divisome proteins is well described in several bacteria, including *Msm* (32). Localization of FtsZ determines the location of the septum, followed by recruitment of PBPs involved in septal PG synthesis and finally enzymes that aid in daughter cell separation (17). Our data shows the temporal dynamics of FtsZ and PBP-lipo septal localization as well as highlights cells where septal co-localization of both proteins can be appreciated.

### Knockdown of PBP-lipo disrupts FtsZ regulation and localization

PBP-lipo repression leads to elongated and branched cells with multiple sites of septation as demonstrated by FDAA staining (Figure 4A). This indicates that PBP-lipo repression disrupts the regulation of cell division in *Mab*. We hypothesized this could be due to dysregulated formation of the FtsZ rings responsible for initiating division. To test this hypothesis, we visualized FtsZ in the context of native PBP-lipo knockdown. We used an N-terminal GFP-FtsZ fusion protein expressed off the FtsZ natural promoter to best capture the natural dynamics of expression. Upon PBP-lipo knockdown, we identified cells that formed multiple FtsZ puncta in contrast to the single puncta found in normally growing cells (Figure 5D). These multiple FtsZ puncta were distributed across the length of the cell indicating that repression of PBP-lipo disrupts FtsZ localization. Furthermore, the formation of multiple FtsZ rings likely leads to the creation of multiple septa and eventually the ectopic branching events observed when PBP-lipo is repressed.

### PBP-lipo is incorporated into a unique PG network in *Mab* that does not exist in *Msm*

Given that *Mtb* and *Msm* do not have duplicates of PBP-lipo in their genomes, we hypothesized that there is a functional difference between PBP-lipo in *Mab* and its homologs in *Msm* and *Mtb*. This possibility intrigued us because an alignment of PBP-lipo from various mycobacteria revealed that PBP-lipo_Mab_ has 5 additional amino acids present only in the *Mab* homolog (Supplemental Figure 9A). When *in silico* structures of PBP-lipo_Mab_ and PBP-lipo_Mtb_ are modeled, these 5 extra residues form an appendage we hypothesized could impart PBP-lipo_Mab_ with additional ability to bind proteins (Supplemental Figure 9B).

To test this hypothesis, we constitutively expressed PBP-lipo_Msm_ in the 2 sgRNA *Mab* strain (hereafter referred to as the *Msm* complement strain). As expected, knockdown of the native *Mab* PBP-lipo impairs growth. However, PBP-lipo_Msm_ expression in the 2 sgRNA strain rescued *Mab* growth (Figure 6A). Furthermore, expressing PBP-lipo_Msm_ eliminated the branching phenotype of the native PBP-lipo knockdown and partially reversed the elongated phenotype as well (Figure 6B). The recoded version of PBP-lipo_Mab_ also reversed the branching of cells and partially complemented the elongated phenotype (Figure 6B). In contrast, the catalytically-inactive version of PBP-lipo_Mab_-(S364A) did not complement, with cells remaining elongated and branched (Figure 6B, Supplemental Figure 10). These results demonstrate that PBP-lipo_Msm_ can rescue knockdown of the endogenous *Mab* enzyme. From these data, we conclude that the essentiality of PBP-lipo_Mab_ is not solely due to a unique functional property that is present only in the *Mab* homolog.

**Figure 6.**
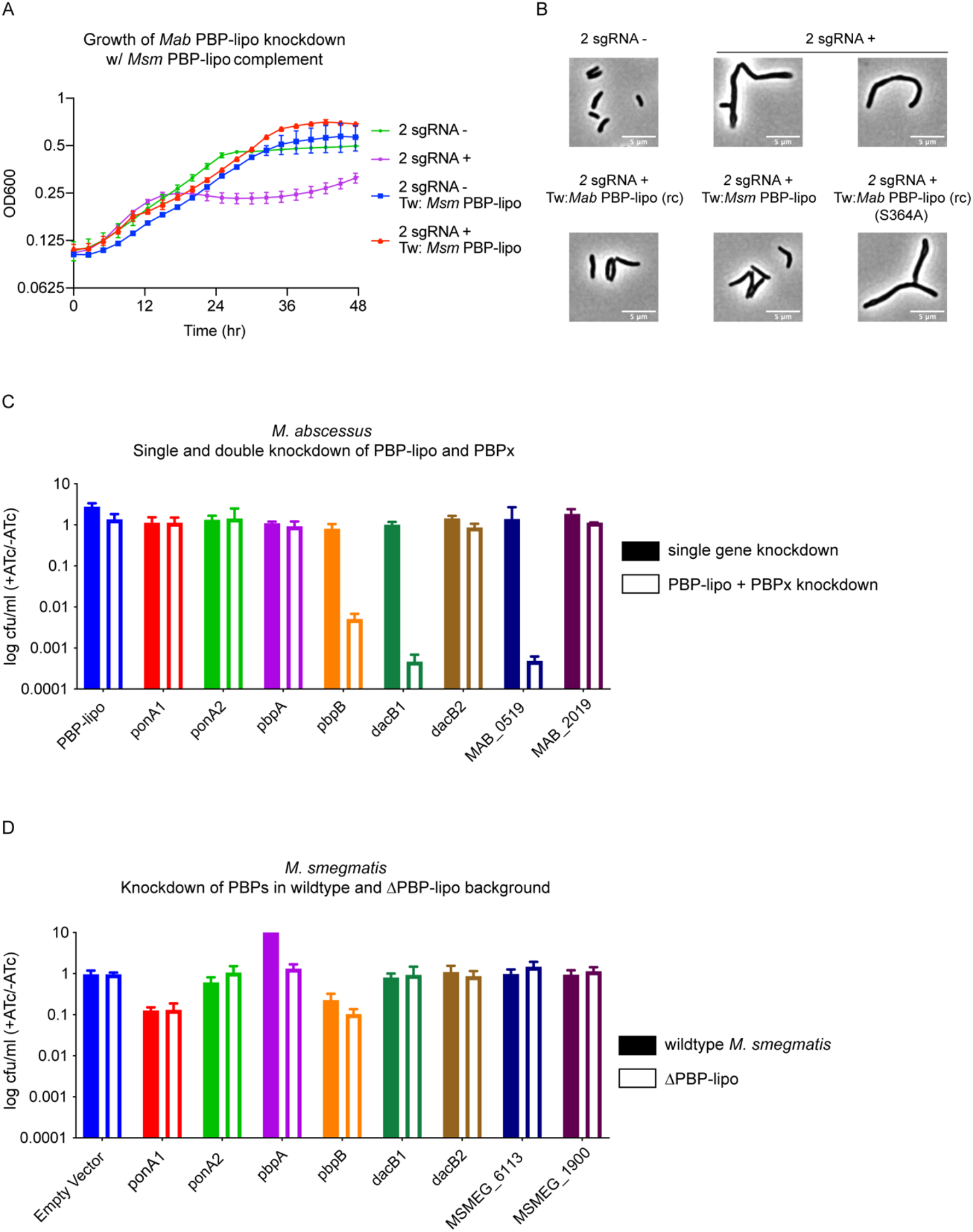
PBP-lipo network in *Mab* does not exist in *Msm*. **(A)** Growth curve of 2 sgRNA strain targeting PBP-lipo and 2 sgRNA strain constitutively expressing PBP-lipo_Msm_. **(B)** Microscopy images of 2 sgRNA and PBP-lipo complement strains. **(C)** Genetic synergy of PBP-lipo with other PBPs in *Mab*. Solid bar represents CFU of strains where a single PBP was knocked down. Open bar represents strains where PBP-lipo was knocked down in combination with the listed PBP. **(D)** Genetic synergy of PBPs and PBP-lipo in *Msm.* CRISPRi plasmids carrying sgRNAs targeting each of the PBPs in *Msm* were transformed into wildtype *Msm* (closed bar) and the ΔPBP-lipo mutant (open bar).

An alternative hypothesis is PBP-lipo participates in a PBP interaction network in *Mab* that is distinct from respective PBP networks in *Msm* and *Mtb*. In short, while PBP-lipo is required in the *Mab* PBP network, its function is likely redundant in the networks of *Msm* and *Mtb.* This notion of functionally distinct PBP networks has been previously described in mycobacteria (36). To interrogate the PBP network of *Mab*, we tested which PBPs genetically interacted with PBP-lipo. To perform this experiment, we cloned CRISPRi constructs with sgRNAs that would weakly knockdown each PBP in *Mab* (PBPx), so that growth was not significantly impaired. We also cloned combinatorial CRISPRi constructs where PBP-lipo and another PBP were weakly repressed. Using this dual sgRNA vector, we could simultaneously knockdown PBP-lipo and a PBP of interest.

Using combinatorial knockdowns of each PBP, we compared growth inhibition when a single PBP was knocked down versus when PBP-lipo and a second PBP were jointly repressed. Of the 8 combinations we tested, we found that knockdown of three PBPs with PBP-lipo lead to synergistic growth arrest (Figure 6C). These genes were *pbpB, dacB1,* and *MAB_0519,* which encodes a protein with a hypothesized carboxypeptidase domain. This genetic synergy was also observed in liquid culture for *pbpB* and *dacB1* (Supplemental Figure 11A). The homolog of PbpB is a well-studied enzyme in both *Msm, Mtb,* and the model organisms *E. coli* and *B. subtilis,* where it is known as FtsI. In all these organisms, PbpB localizes to the septum and is largely responsible for septal PG synthesis (37–39). Interestingly, overexpression of PbpB rescued the growth defect caused by knockdown of PBP-lipo (Supplemental Figure 11B). Similarly, exogenous expression of PbpB fully reversed the morphological defects of PBP-lipo repression (Supplemental Figure 11C). These data indicate that PBP-lipo and PbpB have overlapping functions at the septum in *Mab*. However, because both enzymes are individually essential, PbpB and PBP-lipo must have distinct functions as well.

To test the importance of the genetic interaction of DacB1 with PBP-lipo, we created a DacB1-GFP fusion, which we validated as being fully intact by Western Blot (Supplemental Figure 12A). Interestingly, like PbpB and PBP-lipo, DacB1 localizes to the septum (Supplemental Figure 12B). We also observed co-localization of DacB1 and PBP-lipo at the septal region of the cell (Supplemental Figure 12C). Using the MOMIA and GEMATRIA programs, we quantified the fluorescent signal from DacB1 and PBP-lipo at the septum and binned the cells by cell length. These analyses demonstrate that both DacB1 and PBP-lipo localize to the septum as cells elongate, with DacB1 arriving slightly before PBP-lipo to the septum (Supplemental Figure 12D). These data suggest that PBP-lipo genetically interacts with PbpB and DacB1, and that all three enzymes localize to the same region of the cell and could potentially coordinate septal PG synthesis in a complex together.

The difference in essentiality of PBP-lipo in different species could be due to altered genetic interactions. To test this in *Msm,* we used CRISPRi to knockdown *Msm* PBP homologues in the wildtype *Msm* and the PBP-lipo knockout background. In this experiment, genes that interacted with PBP-lipo would show impaired cell growth when repressed in the PBP-lipo knockout. Interestingly, we did not observe genetic interactions between any of the PBPs tested and PBP-lipo (Figure 6D). Importantly, the level of knockdown for genes that showed synergy in *Mab* was comparable to the level of knockdown in the *Msm* PBP-lipo knockout, as measured by qPCR (Supplemental Figure 13A). We also did not detect synergy in additional PBPs present in the *Msm* genome with no known homologs in *Mab* (Supplemental Figure 13B).

These results might suggest that *Msm* PBP-lipo might have different interacting partners. To test this, we generated high density transposon mutant libraries in both the wildtype and PBP-lipo knockout strains of *Msm* to identify genes that were conditionally essential in the knockout background. The saturation of the 2 wildtype libraries were 51.9% and 53.4%, while the saturation of the PBP-lipo knockout library was 62.4%. Using the resampling method in TRANSIT, our TnSeq data revealed that there were no conditionally essential genes in the ΔPBP-lipo *Msm* background (Supplemental Table 4). These data strongly suggest PBP-lipo in *Msm* exists in a robust genetic network with multiple functional redundancies. To uncover PBP-lipo’s function in *Msm*, it is possible that double and triple knockout mutants of PG synthesis enzymes would need to be generated.

### Repressing PBP-lipo sensitizes *Mab* to a wide range of antibiotics

Given PBP-lipo’s essentiality in *Mab* and the genetic interactions it has with other PBPs, we hypothesized that repressing PBP-lipo would affect *Mab*’s sensitivity to antibiotics, including β-lactams. To test this hypothesis, we measured the sensitivity of PBP-lipo knockdown cells to 21 antibiotics that target translation, transcription, and DNA replication in both *Mab* and *Msm*.

Wildtype *Mab* is typically resistant to β-lactams amoxicillin and ampicillin with MICs greater than >256 µg/ml. When PBP-lipo is knocked down, the MIC for both antibiotics dropped to 2 µg/ml, indicating a greater than 128-fold sensitization (Table 1). These results indicate a strong chemical-genetic interaction between PBP-lipo and both ampicillin and amoxicillin. Interestingly, not all β-lactams had shifts in their MIC when PBP-lipo was knocked down. This suggests that there is a specific chemical-genetic interaction between PBP-lipo or its partner PBPs and ampicillin and amoxicillin. MIC changes for other β-lactams were less drastic, with a 4-8-fold reduction observed for faropenem and cefoxitin (Table 1). PBP-lipo knockdown also affects susceptibility to antibiotics that do not target the cell wall. The macrolides, clarithromycin and erythromycin, were 128-fold and 64-fold more potent respectively in the PBP-lipo knock down cells. A 16-fold reduction in MIC was also observed for RNA polymerase inhibitor, rifampicin.

**Table 1:** Minimum Inhibitory Concentration (MIC) (µg/ml) of ATCC 19977 +/- PBP-lipo Knockdown.

| **Antibiotic** | **-ATc** | **+ATc** | **Fold Difference in MIC**<br>**(-ATc/+ATc)** |
| --- | --- | --- | --- |
| **Cell Wall** |  |  |  |
| Ampicillin | >256 | 2 | >128X |
| Amoxicillin | >256 | 2 | >128X |
| Faropenem | 8 | 1 | 8X |
| Cefoxitin | 8 | 2 | 4X |
| Vancomycin | 8 | 2 | 4X |
| Meropenem | 128 | 128 | N/A |
| Imipenem | >64 | >64 | " " |
| Ticarcillin | >64 | >64 | " " |
| Ceftazidime | >64 | >64 | " " |
| Cephalexin | >64 | >64 | " " |
| Ceftazidime | >64 | >64 | " " |
| D-Cycloserine | >64 | >64 | " " |
| Ethambutol | >64 | >64 | " " |
| Isoniazid | >512 | >512 | " " |
| **Ribosome** |  |  |  |
| Clarithromycin | 16 | 0.125 | 128X |
| Erythromycin | 32 | 0.5 | 64X |
| Amikacin | 16 | 4 | 4X |
| Clindamycin | >64 | 64 | N/A |
| **RNA Polymerase** |  |  |  |
| Rifampicin | 32 | 2 | 16X |
| **DNA Gyrase** |  |  |  |
| Ofloxacin | 8 | 2 | 4X |
| **Other** |  |  |  |
| Pyrazinamide | >64 | >64 | N/A |
| Pretomanid | >64 | >64 | " " |

The MIC differences in rifampicin and clarithromycin are notable because these antibiotics do not target the cell wall. Thus, the mechanism of synergy between PBP-lipo repression and these antibiotics likely differs from the synergy with ampicillin and amoxicillin. We hypothesized that PBP-lipo repression might increase the permeability of the cell wall, allowing more drug to enter the cell. To test this hypothesis, we measured the accumulation of calcein into *Mab*, which is one proxy for cell permeability. Calcein accumulated more rapidly in PBP-lipo knockdown cells, indicating that PBP-lipo increased the permeability of *Mab* cells (Supplemental Figure 14). This increased permeability might lead to increased antibiotic access to the cell, thus sensitizing the bacteria to the antibiotics.

When the MICs of the same 21 antibiotics tested in *Mab* were measured in the PBP-lipo knockout in *Msm,* we observed no difference in the MICs (Supplemental Table 5). We also measured the MICs of a subset of these antibiotics in the PBP-lipo knockout in *Mtb* and found no difference in MICs between the wildtype and knockout strain (Supplemental Table 6). These data suggest that PBP-lipo’s unique function in *Mab’s* PG synthesis network likely contributes to the species’ sensitivity to antibiotics when the enzyme is repressed.

### Repression of PBP-lipo sensitizes *Mab* clinical isolates to antibiotics

The antibiotic experiments were conducted on the lab-adapted ATCC 19977 reference strain on *Mab*, after being isolated from a knee abscess in the 1950s and propagated in the lab since (3, 40). Work performed in *Mtb* has demonstrated clinical isolates and lab-adapted strains respond to antibiotics stressors differently (41). As a result, we tested whether knockdown of PBP-lipo in *Mab* clinical isolates led to the same changes in antibiotic sensitivity as observed in the reference strain.

We found that knockdown of PBP-lipo using the 3 sgRNA construct impaired cell growth in all 11 clinical isolates leading to a 100 to 10,000-fold difference in CFUs (Supplemental Figure 15A). We also confirmed that in a sub-set of the clinical isolates PBP-lipo knockdown lead to elongated and inappropriately branched cells, as observed in the reference strain (Supplemental Figure 15B). After confirming the PBP-lipo phenotypes in the clinical isolates, we measured the MICs of ampicillin, amoxicillin, meropenem, rifampicin, and clarithromycin on PBP-lipo knockdowns of the clinical isolates.

For amoxicillin and ampicillin, PBP-lipo knockdown reduced the MICs by >32 to >512-fold. Meropenem served as a negative control and indeed, there was no difference in MICs between uninduced and induced strains (Table 2). Finally, we observed an 8-32X fold difference in the MICs for rifampicin and 8-128X fold difference in the MICs for clarithromycin (Table 3). Altogether, the antibiotic sensitivity data demonstrate that the phenotypic consequences of repressing PBP-lipo is not just limited to the lab-adapted reference strain, but is observed in *Mab* clinical isolates as well.

**Table 2:** MIC (µg/ml) of *Mab* Clinical Isolates +/- PBP-lipo knockdown (Cell Wall Antibiotics)

| Clinical Isolate | Ampicillin |  |  | Amoxicillin |  |  | Meropenem |  |
| --- | --- | --- | --- | --- | --- | --- | --- | --- |
|  | -ATc | +ATc |  | -ATc | +ATc |  | -ATc | +ATc |
| ATCC 19977 | >256 | 2 |  | >256 | 2 |  | 128 | 128 |
| T35 | >256 | < 0.5 |  | >256 | < 0.5 |  | >256 | 256 |
| T37 | >256 | < 0.5 |  | >256 | < 0.5 |  | >256 | 256 |
| T49 | >256 | 4 |  | >256 | 4 |  | >256 | 128 |
| T50 | >256 | 4 |  | >256 | 4 |  | >256 | 128 |
| T51 | >256 | 2 |  | >256 | 2 |  | >256 | 256 |
| T53 | >256 | 4 |  | >256 | 2 |  | >256 | 256 |
| T56 | >256 | <0.5 |  | >256 | <0.5 |  | >256 | 128 |
| BWH-B | >256 | 8 |  | >256 | 2 |  | >256 | 256 |
| BWH-C | >256 | 4 |  | >256 | 8 |  | >256 | 256 |
| BWH-E | >256 | 4 |  | >256 | 2 |  | >256 | 256 |

**Table 3:** MIC (µg/ml) of *Mab* Clinical Isolates +/- PBP-lipo Knockdown (Rifampicin and Clarithromycin)

|  | Rifampicin |  |  | Clarithromycin |  |
| --- | --- | --- | --- | --- | --- |
| Clinical Isolate | -ATc | +ATc |  | -ATc | +ATc |
| ATCC 19977 | 32 | 2 |  | 16 | 0.125 |
| T35 | 4 | 0.25 |  | 2 | 0.25 |
| T37 | 16 | 1 |  | 2 | 0.25 |
| T49 | 64 | 4 |  | 2 | 0.25 |
| T50 | 32 | 1 |  | 16 | 0.25 |
| T51 | 64 | 4 |  | 16 | 0.5 |
| T53 | 32 | 1 |  | 8 | 0.5 |
| T56 | 8 | 1 |  | 2 | 0.25 |
| BWH-B | 32 | 1 |  | 16 | 0.25 |
| BWH-C | 32 | 2 |  | 32 | 1 |
| BWH-E | 16 | 1 |  | 16 | 0.125 |

## Discussion

Developing new drugs and drug regimens against *Mab* requires a better understanding of the bacterium’s essential processes. This includes identifying essential *Mab* genes for further characterization as potential drug targets. To do this, we generated high-density transposon libraries to identify essential genes in the reference strain, *M. abscessus subsp. abscessus* ATCC 19977. Importantly, *Mab* transposon libraries with fewer than 10,000 unique mutants have been previously published, but these studies did not identify the full list of essential genes in *Mab* (10, 11). Recently, Rifat *et al* successfully generated high-density libraries and were the first group to identify essential genes and genomic elements in the *Mab* genome (12). All together, they identified 326 essential genes and compared this list to essential genes identified in *Mtb* and *Mycobacterium avium.* Similar to our study, Rifat *et al* highlighted uniquely essential genes in *Mab,* which they also state deserve further study as these genes could represent *Mab* specific drug targets (12).

In addition to identifying essential genes in *Mab*, our study builds upon Rifat *et al’s* work by using CRISPRi to repress gene expression and validate predicted essential genes as required for *in vitro* growth (24). Furthermore, we found that by targeting genes with multiple sgRNAs, CRISPRi can achieve significant gene repression, allowing for the investigation of essential genes and providing an alternative to gene knockouts. Constructing gene knockouts in *Mab* can be inefficient (42); thus, CRISPRi provides a convenient tool for genetic manipulation in *Mab*.

Overall, the newly generated *Mab* transposon libraries can be used to determine genes required for growth in a variety of conditions including infection models, biofilms, and antibiotic stress. Both Rifat *et al’s* and our TnSeq data identified PBP-lipo as the sole PBP that is essential in *Mab* and non-essential in *Mtb* (12). Using CRISPRi, we demonstrated the necessity of PBP-lipo expression for *Mab* to maintain normal morphology and to divide properly. When PBP-lipo is repressed, cells elongate, branch, and form ectopic poles. We localized PBP-lipo to the subpolar regions of the cell, and as cells elongate and prepare for division, PBP-lipo localizes to the septum. Furthermore, depleting PBP-lipo led to the creation of multiple Z-rings in the cell and likely the aberrant start of cell division and formation of branched cells Overall, the constellation of PBP-lipo knockdown phenotypes mimic septal-factor depletions observed in *Msm,* which also cause cells to elongate, branch, and form ectopic poles (32). These data suggest that PBP-lipo is involved with PG synthesis at the septum and that depletion of the enzyme leads to dysregulated septal PG synthesis and cell division.

Given PBP-lipo’s localization and effect on cell division, we propose that PBP-lipo acts as an essential septal PG synthesis enzyme in *Mab.* In *Mtb* and *Msm,* this is accomplished by the enzyme *pbpB*, which is essential for growth. Additionally, when *pbpB* is repressed in these species, cells appear morphologically similar to the PBP-lipo knockdown in *Mab* (32, 37). Interestingly, PBP-lipo knockouts in both *Msm* and *Mtb* are morphologically identical to wildtype cells (Supplemental Figure 7). These data underscore that biological insights from *Msm* and *Mtb* should not be de-facto applied to *Mab*. We hypothesize that PBP-lipo is a member of the *Mab* divisome complex, a collection of structural proteins and enzymes that coordinates cell division (43). This hypothesis was supported by the co-localization of PBP-lipo with FtsZ, a key initiating component of the bacterial divisome (34). Future studies are needed to determine if PBP-lipo interacts with known septal factors, such as FtsQ and FtsW, whose functions in the divisome have been described in *Msm* (44–46). Future work may also reveal novel PBP-lipo binding factors that coordinate PG synthesis and cell division in *Mab* specifically.

Phenotypes characteristic of the PBP-lipo knockdown in *Mab* were absent in the *Mtb* and *Msm* PBP-lipo knockouts. Both *Mtb* and *Msm* do not have a duplicate copy of PBP-lipo in their genomes. Thus, to explain the differential essentiality of PBP-lipo across these mycobacterial species, one hypothesis we explored was differences in the genetic networks of PG synthesis enzymes. Construction of the PG layer is a complex process that involves a collection of enzymes that work in concert to ensure proper synthesis and remodeling. Previous work has shown that PG synthesis enzymes belong to genetic and spatiotemporal networks that do not necessarily replicate across closely related species (36, 47). Thus, while the synthetic enzymes to construct the PG layer may be homologous between species, how the enzymes are networked and subsequently function together may differ. We demonstrated that PBP-lipo genetically interacts with *pbpB, dacB1,* and *MAB_0519*. In contrast, TnSeq on the *Msm* Δ*pbp-lipo* strain did not reveal interactions with any other gene in the genome. This demonstrates that despite being homologs, PBP-lipo in *Mab* and *Msm* are incorporated into different PG synthesis networks. In *Mab,* this network renders PBP-lipo’s function as essential for growth and proper cell division, whereas in *Msm,* PBP-lipo is dispensable. Interestingly, in *Mtb* PBP-lipo is required for normal growth in a *ponA2* knockout suggesting that PBP-lipo and PonA2 genetically interact *Mtb* (36) *–* a genetic interaction that was not observed in *Mab*. This observation supports the hypothesis that PBP-lipo is incorporated in different PG synthesis in *Mtb* as well.

While this work demonstrates the essential function of PBP-lipo for growth and division in *Mab* and not in *Msm* or *Mtb,* all the experiments were performed in standard growth conditions. Studies in other bacteria have demonstrated that homologous PBPs that are otherwise non-essential become required under specific growth conditions. For example, in *Salmonella*, the class B homolog of PBP3, PBP3-SAL, was required for cell division in acidified intraphagosomal environments (48). Similarly, in *E. coli*, PBP6, which encodes a carboxypeptidase, was more active at lower pH values compared to 5 other homologous carboxypeptidases (49). These studies demonstrate that some PBPs become more active and their functions more required in specific growth conditions. Future studies could test PBP-lipo’s function under various stress conditions. These experiments may elucidate a condition-specific role for PBP-lipo in *Msm* and *Mtb* that differs from its central function in *Mab*.

Not only is PBP-lipo essential in *Mab,* it also genetically interacts with three other PBPs, *pbpB, dacB1,* and *MAB_0519,* which are all hypothesized to be septal associated. PbpB is a D,D-transpeptidase that performs 4,3 crosslinking reactions at the septum. In *Msm*, depleting PbpB leads to cell filamentation and branching reminiscent of the *Mab* PBP-lipo knockdown phenotypes (32). A potential explanation of this genetic interaction is both PbpB and PBP-lipo perform 4,3-crosslinking at the *Mab* septum. In this model, repressing both enzymes would dramatically reduce 4,3-crosslinking at the septum and lead to cell death. We explored this model by overexpressing PbpB while strongly repressing PBP-lipo. We discovered overexpressing PbpB completely reversed the growth and morphological defect of PBP-lipo knockdown. This data provided evidence that PBP-lipo and PbpB have overlapping 4-3 cross-linking function at the septum, and that by supplying the cell with more D,D-transpeptidation activity via PbpB, the deleterious effects of PBP-lipo repression were reversed. Interestingly, given that both PBP-lipo and PbpB are essential in *Mab*, the enzymes must have overlapping, but distinct functional roles. Their distinct functions could be revealed by identifying unique genetic or physical interactions each enzyme has with other PG synthesis enzymes. Why *Mab* requires two PG 4,3 crosslinking enzymes functioning at the septum is an intriguing area of future research.

The two other genes that synergized with PBP-lipo, DacB1 and MAB_0519, are both hypothesized carboxypeptidases that cleave the terminal D-ala-D-ala on the pentapeptide chains of PG monomers forming a tetrapeptide. This substrate is necessary for 3,3 crosslinks catalyzed by L-D transpeptidases (LDTs) (50, 51). Interestingly, DacB1 localizes to the septum in *Mab* and has been described as septally localized in *Msm* and *Mtb* (52). In *Mab,* DacB1 also co-localizes with PBP-lipo at the septum. Furthermore, the homolog of MAB_0519 in *Mtb*, Rv3627c is involved in septal PG synthesis (53). Thus, we hypothesize DacB1 and MAB_0519 are involved in septal PG synthesis and may work with PBP-lipo to ensure proper PG synthesis and remodeling at the septum. Interestingly, DacB1 and MAB_0519 are not functionally redundant, given that both genes are synergistic with PBP-lipo. Further characterization of the PG enzymes in *Mab* and their genetic networks will uncover species-specific nuances in *Mab* PG synthesis that have the potential to be exploited for drug discovery.

Knockdown of PBP-lipo dramatically sensitized the reference *Mab* strain and 11 clinical isolates to several antibiotics. The PBP-lipo knockdown was >128-fold more sensitive to the β-lactams, ampicillin and amoxicillin, while other β-lactams such as like meropenem, cephalexin, ticarcillin and others showed no difference in MIC. This result indicates a specific chemical-genetic interaction between ampicillin/amoxicillin, and PBP-lipo. Historically, β-lactams have not been used to treat *Mab* infections due to high levels of resistance and the presence of β-lactamases (20). While imipenem and cefoxitin are used with some success, macrolides and aminoglycosides remain the backbone of *Mab* treatment (54). Recently, studies have shown dual β-lactam strategies are effective in killing *Mab* both *in vitro* and *in vivo* (55–57). These synergy data suggest that understanding the specific antibiotic targets of β-lactams in *Mab* would augment the design of effective dual β-lactam combinations. At the moment, effective combinations are usually tested empirically with no insight into potential mechanisms (56). Recently, a potential mechanism for dual-beta lactam synergy in *Mab* was uncovered by Dousa *et al.* The group discovered that the novel beta-lactamase inhibitor, durlobactam, also inhibited a *Mab* carboxypeptidase and by doing so, greatly enhanced *Mab* sensitivity to amoxicillin and imipenem combination therapy (58). Another attempt to identify specific targets of PBP targets of beta-lactams was conducted by Sayed *et al*. This group used a combination of a bocillin FL binding assay to stain all the PBPs in *Mab* followed by proteomics to identify the targets of 12 β-lactams and 2 β-lactamases. Interestingly, in their datasets, PBP-lipo was not identified by mass spectrometry of bocillin FL-bounded PBPs (59). This further supports our hypothesis that PBP-lipo is expressed at low-levels in *Mab* and is not identifiable by bocillin FL staining under native expression conditions.

Nonetheless, our data demonstrate that using a β-lactam or small molecules that target PBP-lipo would not only kill the cell, but also potentiate β-lactams like ampicillin and amoxicillin. In addition, the permeabilizing effect of PBP-lipo inhibition would potentiate drugs by allowing more compound to enter the cell. Crucially, the observed synergy with PBP-lipo knockdown and β-lactams was not only limited to the reference strain, but also present in 11 clinical isolates of *Mab,* indicating that this synergy can be broadly targeted in *Mab* clinical isolates.

Overall, this work employed TnSeq to identify the uniquely essential gene PBP-lipo in *Mab*. We present PBP-lipo as a promising drug target given both its essentiality in *Mab* cell growth and division and its role in sensitizing *Mab* to a range of antibiotics, including commonly used and accessible antibiotics, ampicillin and amoxicillin. Future studies exploring the PBP networks of mycobacteria can expose further species-specific vulnerabilities in peptidoglycan synthesis. Finally, identifying chemical inhibitors of PBP-lipo may help form the foundation for new treatments against this emerging and difficult to treat pathogen.

## Materials and Methods

A full list of the strains, plasmids, and primers is available in Supplemental Tables 7, 8, and 9 respectively.

### Bacterial Strains and Culture Conditions

All *Mab* and *Mtb* strains were grown shaking at 37°C in liquid 7H9 media consisting of Middlebrook 7H9 salts with 0.2% glycerol, 0.85 g/L NaCl, OADC (oleic acid, 5 g/L albumin, 2 g/L dextrose, 0.003 g/L catalase), and 0.05% Tween80. *Mab* and *Mtb* were plated on Middlebrook 7H10 agar supplemented with 0.5% glycerol. *Msm* was grown shaking at 37°C in liquid on 7H9 media consisting of Middlebrook 7H9 salts with 0.2% glycerol, 0.85 g/L NaCl, ADC, and 0.05% Tween80 and plated on LB agar. Antibiotic selection concentrations for *Mab* cultures were kanamycin 50 µg/µl and zeocin 100 µg/µl. For transposon mutant selection, 100 µg/ml of kanamycin was used. Antibiotic selection concentrations for *Msm* and *Mtb* cultures were kanamycin 25 µg/µl, zeocin 20 µg/µl, and gentamicin 5 µg/µl. For *E. coli* (TOP10, XL1-Blue and DH5α), all antibiotic concentrations were twice those of the concentrations used for *Msm* and *Mtb*. Induction of all CRISPRi plasmids was performed with 500 ng/ul anhydrous tetracycline (ATc).

### Strain Construction

*Construction of 0, 1, 2, 3 sgRNA PBP-lipo knockdown strains and PBP-lipo complement strains* To build the 0 sgRNA (empty sgRNA) *Mab* strain, we transformed the pCT296 vector containing an empty sgRNA into *Mab.* To build the 1, 2, and 3 sgRNA strains, first we built individual CRISPRi plasmids containing single sgRNAs that targeted PBP-lipo. These plasmids were cr25, cr26, and cr56. We then designed primers that amplified the sgRNA targeting region with the appropriate flanking sequences and performed Golden Gate cloning to combine the inserts into one vector. This method was previously described by Rock et al (24). For the 1 sgRNA strain, we transformed cr25 into the *Mab* strain. For the 2 sgRNA, we transformed the vector pCRC1, which contained the cr25 and cr26 sgRNAs on one backbone. Finally, for the 3 sgRNA strain we transformed pCRC2, which contained the cr25, cr26, and cr56 sgRNAs on one backbone. To build the PBP-lipo complement strain, we designed primers to amplify PBP-lipo with modified protospacer adjacent motif (PAM) sequences that were no longer homologous to the 1 and 2 sgRNAs. This recoded PBP-lipo was then cloned into a constitutive expression vector resulting in the pCA323 construct, which was transformed into the 2 sgRNA strain. The catalytically dead mutant version of the recoded PBP-lipo was generated by using primers to generate the S364A mutation. This engineered PBP-lipo was cloned into a constitutive expression vector and the resulting construct, pCA324 was transformed into the 2 sgRNA strain.

#### *Msm* and *Mtb* ΔPBP-lipo strains

To generate PBP-lipo knockout mutants in *Msm* and *Mtb,* we used a recombineering approach to replace the endogenous copy of the gene with a zeocin resistance cassette as previously described (60). First, 500 base pairs of upstream and downstream sequence surrounding PBP-lipo (*Rv2864c* and *MSMEG_2584c*) were amplified by PCR KOD XtremeTM Hot Start DNA polymerase (EMD Millipore, Billerica, MA). These flanking regions were amplified with overlaps for the zeocin resistance cassette. The 2 flanking regions and zeocin resistance cassette were then assembled via isothermal assembly (61) into a single construct, which was then amplified by PCR. Each deletion construct was transformed into either *Mtb* or *Msm* expressing inducible copies of RecET (62). The resulting colonies were screened for the PBP-lipo deletion by PCR.

#### PBP-lipo_Mab_-strep expression strain

PBP-lipo_Mab_ was amplified using primers that added a C-terminal strep tag to the protein. The resulting insert was Gibson stitched to the vector backbone that drove expression of PBP-lipo_Mab_-strep from the constitutively active MOP promoter.

#### mRFP-PBP-lipo_Mab_

We amplified PBP-lipo_Mab_ in two fragments. The first fragment contained the N-terminal signal sequence of PBP-lipo. The second fragment contained the remainder of the protein. We then amplified mRFP with overhangs that were compatible with the two fragments of PBP-lipo. The resulting three fragments were Gibson assembled to a vector backbone that constitutively drove expression of the mRFP-PBP-lipo_Mab_ fusion protein. A gly-gly-ser-gly-ser-gly linker was used in between mRFP and the two fragments of PBP-lipo.

#### DacB1-GFP-strep

DacB1 lacking a stop codon was amplified along with GFP containing overlaps with DacB1 and a gly-gly-ser linker. The two fragments were Gibson assembled to a vector backbone that constitutively drove expression of the DacB1-GFP-strep fusion protein.

#### ftsZ-mNeonGreen & DacB1-mNeonGreen

ftsZ and DacB1 were amplified along with mNeonGreen containing a gly-gly-ser linker. Each gene and its corresponding overlapping mNeonGreen fragment were Gibson assembled to a vector that constitutively drove expression from the weak promoter iMyc.

#### natP-GFP-ftsZ

To express FtsZ off its natural promoter, 300bp of sequence upstream of the FtsZ start codon was amplified and Gibson assembled with 2 additional fragments corresponding to GFP with a gly-gly-ser linker and FtsZ. The resulting vector was an N-terminal GFP-FtsZ fusion expressed off the FtsZ natural promoter.

### Growth Curve

*Mab, Msm,* and *Mtb* cultures were grown to log phase and diluted to OD600 0.05. Growth curves were performed in a 96 well format at 37°C with shaking. OD600 was measured every 15 min for 24-72 hr using the TECAN plate reader

### Colony Forming Units

*Mab* and *Msm* cultures were grown to mid-log phase and then serially diluted in their respective media in 96 well plates. Dilutions were then spot plated onto solid media +/- 500 ng/ul ATc. Plates were incubated at 37°C for 4 days *(Mab)* or 3 days *(Msm)*. After incubation, the resulting CFUs were calculated by counting the number of colonies at the appropriate dilution. The fraction survivability was calculated as CFUs +ATc / -ATc.

### Minimum Inhibitory Concentration Determination

*Mab, Msm,* and *Mtb* were grown to mid-log phase and diluted to an OD600 = 0.003 in each well of non-treated 96-well plates (Genesee Scientific) containing 100 μL of antibiotic serially diluted in 7H9 + OADC + 5 μg/mL clavulanate (Sigma Aldrich). For MICs on *Mab* knockdown cells, cultures were induced for knockdown 18 hr prior with 500 ng/ul ATc. Wells with knockdown bacteria also contained 500 ng/ul ATc. *Msm* media contained ADC rather than OADC. Cells were incubated with drug at 37°C with shaking for 1 day *(Mab, Msm)* or 7 days *(Mtb)*. Afterwards, 0.002% resazurin diluted in ddH_2_O (Sigma Aldrich) was added to each well. Plates were then incubated for 24 hr. MICs were determined by the concentration of antibiotic that turned wells blue signifying no metabolic activity (63).

### Generation of Transposon Libraries

To create transposon mutant libraries of *Mab* ATCC 19977 and *Msm* (wildtype and ΔPBP-lipo), 100 ml of bacterial cultures were grown to an OD600 of 1.5 – 2.0. For *Mab*, cultures were grown in biological triplicate. For *Msm*, the wildtype strain was grown in duplicate while one replicate of the ΔPBP-lipo strain was grown. The cells were then washed twice with 50 ml of MP Buffer (50mM Tris-HCl pH 7.5, 150 mM NaCl, 10 mM MgSO_4_, 2 mM CaCl_2_) and resuspended in 1/10th the culture volume in MP Buffer. Afterwards, 2 x 10^11^ pfu of temperature sensitive φMycoMarT7 phage carrying the Himar1 transposon were added to bacteria. Phage and bacterial cultures were incubated at 37°C for 4 hr with shaking. The transduced cultures were spun down at RT 4000 rpm for 10 min and resuspended in 12 ml of PBS-Tween. The cultures were then tittered by plating on 7H10 plates (*Mab)* or LB *(Msm)* supplemented with kanamycin 100 µg/µml *(Mab)* or 20 µg/µl *(Msm)* before freezing the aliquots for future plating. After determining the titer, 150,000 bacterial mutants of each transduced culture were plated onto kanamycin selective plates and grown for 4 days at 37°C. The resulting mutant libraries were harvested and stored in aliquots with 7H9 + 10% glycerol at −80°C.

### Genomic DNA Extraction

To prep libraries for sequencing, we isolated the gDNA of the mutant libraries using a bead-beating protocol developed in the lab. 2-6 ml of transposon mutant libraries were thawed and spun down to remove excess glycerol in the sample. Pellets were then resuspended in TE Buffer (10 mM Tris HCl pH 7.4, 1 mM EDTA pH8) and placed in bead-beating tubes with 600 µl of 25:24:1 phenol:chloroform:isoamyl alcohol. The samples were then bead-beat 4X for 45 s at 4000 rpm. Samples were cooled on ice for 45 s between each successive bead-beat round. After bead-beating, the samples were spun down for 10 min at 13,000 rpm. The supernatant was transferred to a new falcon tube and incubated with 1:1 volume phenol:chloroform for 1 hr rocking. Samples were then spun down in MaXtract High Density phase-lock tubes (Qiagen) at 1500 g for 5 min. This separated the aqueous and organic layers. Afterwards, ½ volume of chloroform was added to the tube and the contents were spun down at 1500 g for 5 min. The aqueous phase was transferred to a new tube and RNAse A (Thermo Fisher) was added to a final concentration of 25 µg/µl. The tubes were incubated at 37°C for 1 hr with shaking. Finally, the supernatant was washed with 1:1 volume of phenol-chloroform followed by ½ volume of chloroform. The aqueous phase was transferred to a new tube and genomic DNA was precipitated with 1 volume of isopropanol alcohol and 1/10th volume of 3 M sodium acetate pH 5.2. The library gDNA was washed twice with fresh 70% ethanol and resuspended in nuclease free ddH_2_O. This protocol consistently generated high yields of *Mab* gDNA.

### Transposon sequencing, mapping, and analysis

Sequencing libraries were prepared from the extracted gDNA by amplifying chromosomal-transposon junctions following the protocol as described by Long et al, 2015 (21). These amplicons were then sequenced using the Illumina Hi-Seq platform. The resulting reads were mapped onto the *Mab* or *Msm* genomes. The data were analyzed using the TRANSIT pipeline (23). Insertion counts at each TA site were normalized using TTR and averaged across replicates. Essentiality categories of genes were determined by the Hidden Markov Model (HMM) algorithm in TRANSIT. Resampling analysis was used to compare insertion accounts between genes in the wildtype and ΔPBP-lipo *Msm* strains.

### Cloning of CRISPRi Constructs

To clone sgRNAs onto the CRISPRi vector backbone, complimentary targeting oligos were annealed and ligated (T4 DNA ligase, NEB) into the BsmBI digested CRISPRi vector backbone. To clone multiple sgRNAs into the same vector, the promoter, sgRNA handle, and sgRNA sequences were amplified with the appropriate flanking sequences to perform Golden Gate cloning using the SapI cloning site in the CRISPRi vector (24). All cloning was performed using DH5α cells and sequences were verified before transformation into *Mab* and *Msm*.

### Transformation of *Mab*

To generate electrocompetent *Mab* cells, 50 ml of *Mab* cultures were grown to mid-log phase. Cells were spun down at 4000 rpm for 10 min and washed twice with pre-chilled 10% glycerol. Cell pellets were resuspended in 1/100th of the starting volume and kept on ice for immediate use or stored at −80°C for future use. For transformations ∼100 ng of plasmid DNA was incubated with electro-competent cells for 5 min before electroporation using the settings: 2500 V, 125 Ω, 25 μF. Afterwards, cells were recovered in 1 ml of 7H10 with OADC without selection for 3 hr at 37°C with shaking. Cells were then plated on 7H10 with the appropriate selection marker and incubated for 4 days at 37°C.

### Microscopy and Image Analysis

All imaging was performed on an inverted Nikon TI-E microscope at 60X and 100X magnification. For PBP-lipo knockdown experiments, cultures were induced for knockdown with 500 ng/ul ATc for 18 hr prior to imaging. All *Mab* cultures were fixed with 7H9 + 3% paraformaldehyde (PFA) for 1 hr and resuspended in PBS + 0.05% Tween80 prior to imaging. Cellular features including cell length, width, and fluorescence signal were analyzed using the MOMIA and GEMATRIA image analysis pipelines developed in the lab (35).

### Fluorescent D-Amino Acid Staining

NADA (3-[(7-Nitro-2,1,3-benzoxadiazol-4-yl)amino]-D-alanine hydrochloride) was synthesized by Tocris following the published protocol (31). To stain *Mab* with NADA, 0.1 mM of FDAA was added to 1 ml of exponentially growing cells and incubated for 3 min before washing in warm 7H9 twice. For *Mab* imaging, after the second wash, cells were fixed in 7H9 + 3% paraformaldehyde for 1 hr. The PFA was then washed off and cells were resuspended in PBS + 0.05% Tween80 prior to imaging.

### RNA Extraction and RT-qPCR

For each strain, cultures were grown in biological triplicate to mid-log phase and then diluted back in +/- 500 ng/ml ATc and grown for 18 hr to achieve target knockdown. Afterwards, 2 OD 600 equivalents of cells from each culture were harvested by centrifugation, resuspended in TRIzol (Thermo Fisher), and lysed by bead beating (Lysing Matrix B, MP Biomedicals). Total RNA was extracted using the RNA miniprep (Zymo Research). Residual genomic DNA was digested with TURBO DNase (Ambion), and samples were cleaned with RNA clean-up columns (Zymo Research). cDNA was prepared using random hexamers following manual instructions (Life Technologies Superscript IV). Alkaline hydrolysis was then used to remove RNA. cDNA was purified by spin column (Qiagen) and then quantified by real-time quantitative PCR (RT–qPCR) on a Viia7 light cycler (Applied Biosystems) using iTaq Universal SYBR Green Supermix (BioRad). All qPCR primer pairs were confirmed to be >95% efficient. The masses of cDNA used were experimentally validated to be within the linear dynamic range of the assay. Signals were normalized to the *sigA* (*MAB_3009*) transcript and quantified by the ΔΔCt method. Error bars are standard deviations from three biological replicates.

### Bocillin FL staining

*Mab* cultures were grown to mid-log phase and 10 OD units of bacteria were spun down and resuspended in 1ml of fresh 7H9 media with 5 µg/ml of clavulanate. Bocillin-BODIPY or FITC were added to cultures to a final concentration of 10 µM. Cells were then incubated at 37°C for 2.5 hr, with aluminum wrapping to prevent light exposure. After labeling, cells are washed with PBS + 0.05% Tween80 three times and resuspended in 200 µl of lysis buffer (50 mM Tris-HCl pH = 7.4, 50 mM NaCl, protease inhibitor (Roche)). Samples were bead beat 4X at 4000 rpm for 1 min with 1 min of rest on ice in between rounds of bead-beats. CaCl_2_ was added to a final concentration of 1 mM along with 4U of DNAse I and 10 mg/ml of fresh lysozyme. Samples were incubated at 37°C for 30 min. Afterwards, supernatants were normalized by A280 protein concentration, diluted with 6X Laemmli buffer, and run on 4%– 12% NuPAGE Bis Tris precast gels (Life Technologies). To visualize fluorescence, we used a BioRad Gel Doc system. To detect the presence of PBP-lipo-strep, we performed a Western Blot. Membranes were blotted with rabbit α-Strep (GenScript) at 1:1000 in TBST + 3% BSA.

### Calcein Staining

For each strain, cultures were grown in biological triplicate to mid-log phase and diluted back in +/- 500 ng/ml ATc and grown for 18 hr to achieve target knockdown. Afterwards, 3 OD units of bacteria were washed twice with PBS + 0.05% Tween80. Cells were resuspended in PBS + 0.05% Tween80 and 100 µl of cells at OD600 = 0.4 was added to 96 well plates. The plates were incubated in the plate reader for 30 min at 37°C with shaking. Afterwards, calcein was added at a final concentration of 1 µg/ml. Fluorescence signal was measured every minute using the Tecan plate reader.

## Acknowledgements

This project was funded by the generous support of the Paul and Daisy Soros Foundation, the NIH/NIAID F31 Predoctoral Fellowship award number F31AI149932, and award Number T32GM007753 from the National Institute of General Medical Sciences. The content is solely the responsibility of the authors and does not represent the official views of the National Institute of General Medical Sciences or the National Institutes of Health.

## Notes

### Competing Interest Statement

The authors have declared no competing interest.

### Summary of Updates

-New data identifying the spatiotemporal dynamics of PBP-lipo's localization relative to FtsZ -Data showing the effect of PBP-lipo knockdown on FtsZ localization -Data showing that the mRFP-PBP-lipo fusion protein is intact and functional -Data showing that overexpression of PbpB can rescue PBP-lipo knockdown -Authors and author affiliations updated.

