## Supplemental Figures for "High-density transposon mutagenesis in *Mycobacterium abscessus* identifies an essential penicillin-binding lipo-protein (PBP-lipo) involved in septal peptidoglycan synthesis and antibiotic sensitivity"

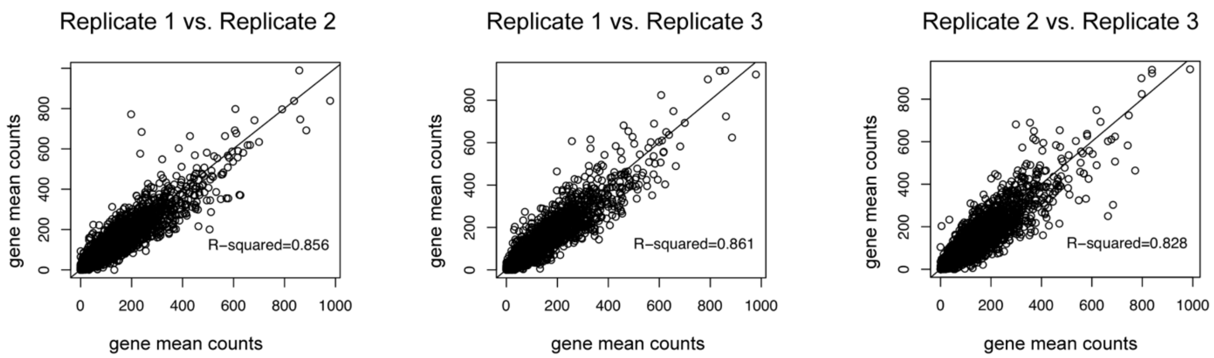


**Supplemental Figure 1. Correlation of *Mab* TnSeq Libraries.** Pairwise comparisons of transposon insertions in each gene between two replicate libraries. Pearson correlation was calculated for each comparison and the $r^2$ value is reported.

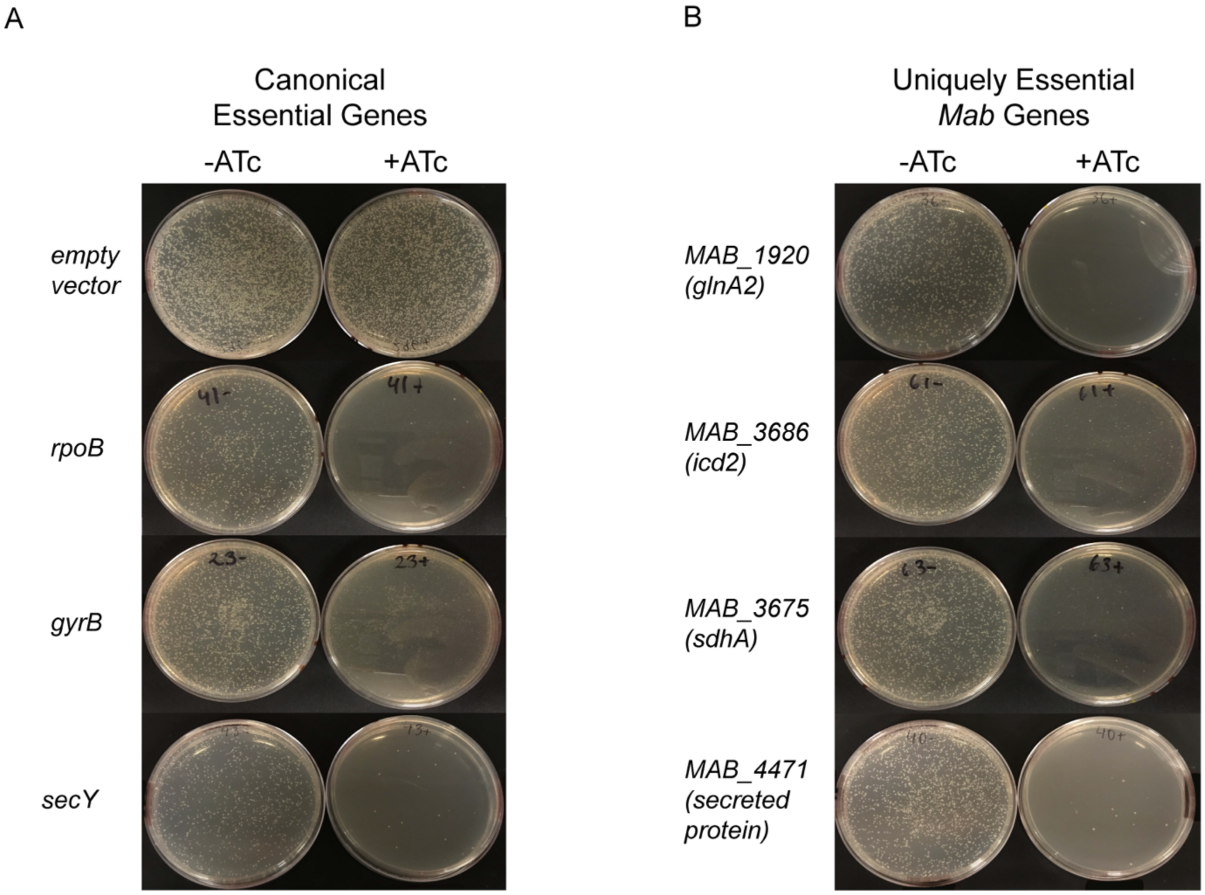


**Supplemental Figure 2**. **Validation of Uniquely Essential *Mab* Genes. (A)** Transformation of CRISPRi plasmids carrying sgRNAs targeting canonical and **(B)** uniquely essential genes in *Mab*. Equal volume of transformations was plated on +/- ATc plates.

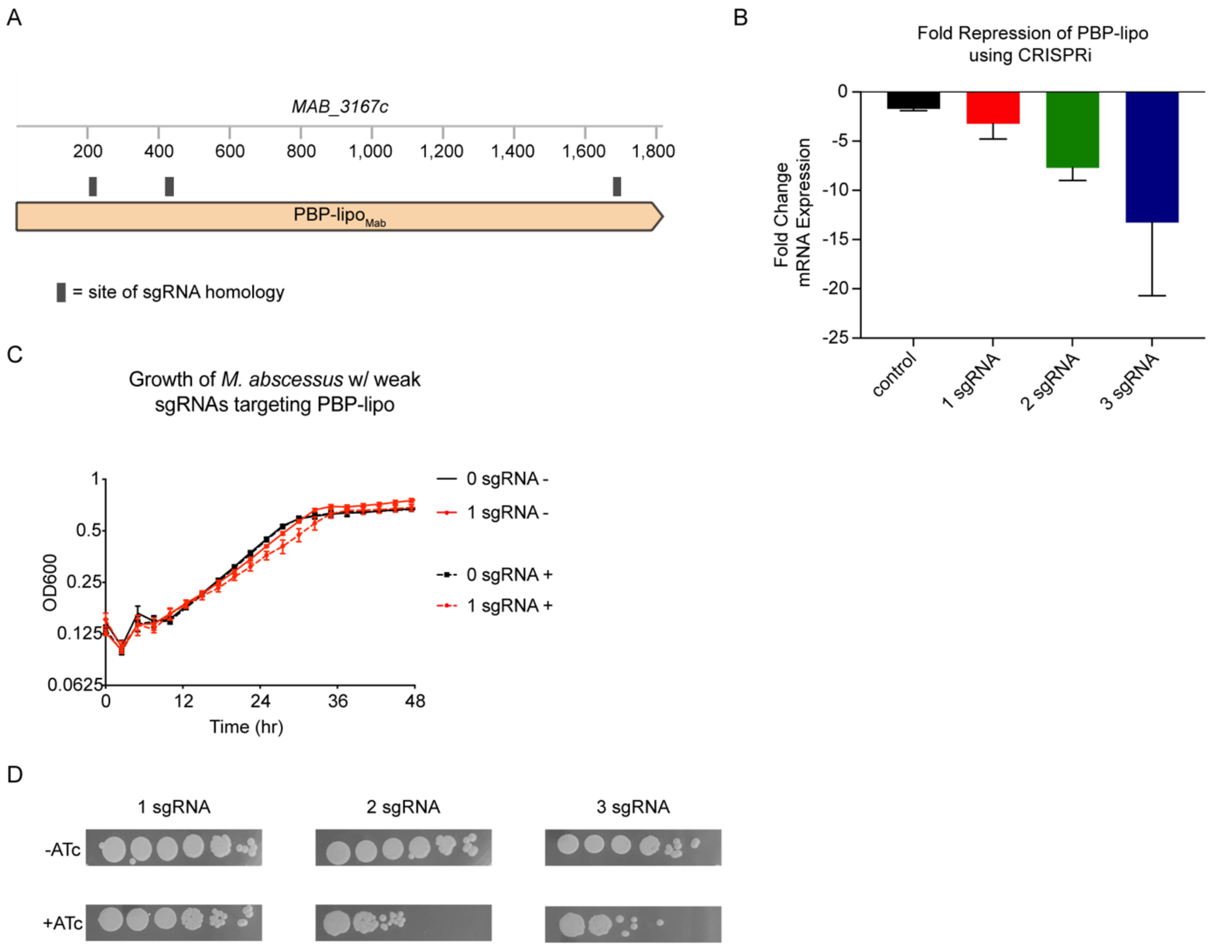


**Supplemental Figure 3. Knockdown of PBP-lipo impairs *Mab* growth. (A)** Schematic of *MAB_3167c*, encoding PBP-lipo. Colored rectangles indicate sgRNAs binding sites **(B)** Fold change in mRNA expression measured by qPCR for empty sgRNA (negative control) and 1, 2, and 3 sgRNAs targeting PBP-lipo. **(C)** Growth of *Mab* cultures transformed with CRISPRi plasmid carrying either 0 sgRNA (empty guide control) or 1 sgRNA targeting PBP-lipo. "-" symbol indicates cultures were grown without ATc. "+" symbol indicates cultures were grown with ATc. **(D)** CFU of *Mab* with 1, 2, or 3 sgRNAs targeting PBP-lipo. Cultures were spotted on 7H10 plates +/- ATc.

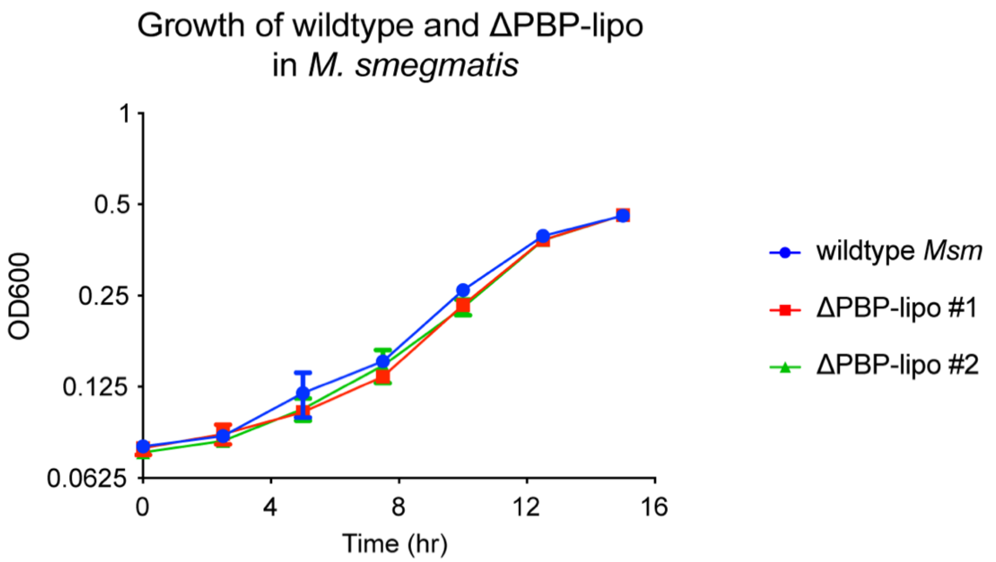


**Supplemental Figure 4. Growth of wildtype *Msm* and PBP-lipo knockout strains.**

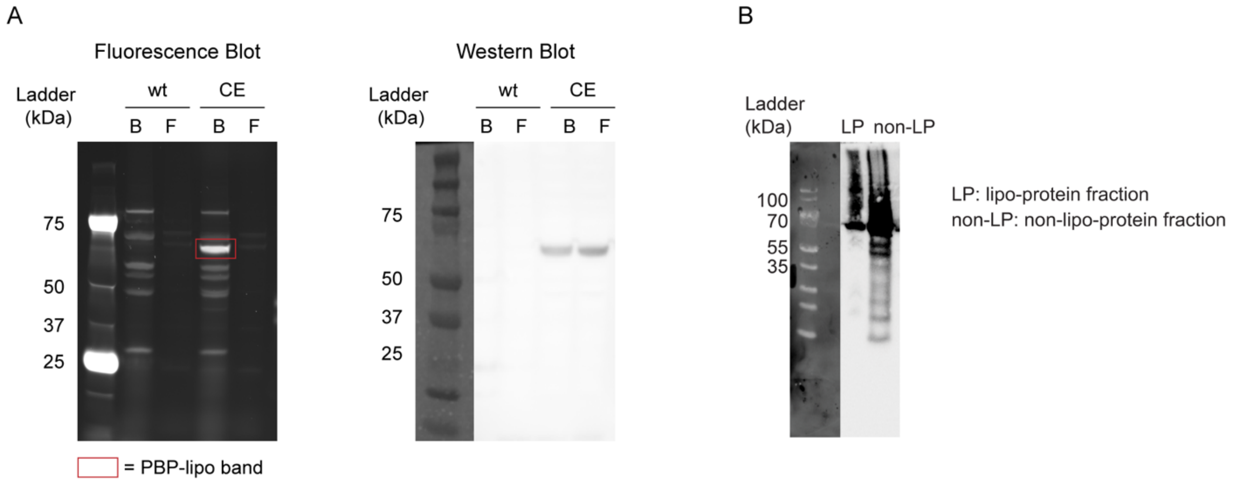


**Supplemental Figure 5. (A) *MAB_3167c* encodes a functional PBP in *Mab*.** Bocillin FL staining of PBP-lipo-strep. Wildtype *Mab* (wt) and a PBP-lipo-strep tagged constitutive strain (CE) were stained with bocillin FL (B), a fluorescent penicillin analog and FITC (F) as a negative control. (*left*) Blot shows PBPs bound to bocillin FL. (*right*) Blot shows α-strep Western blot for PBP-lipo-strep. **(B)** Western Blot of lipoprotein (LP) and non-lipoprotein (non-LP) fractions of *Mab.* PBP-lipo-strep was detected in both fractions.

A

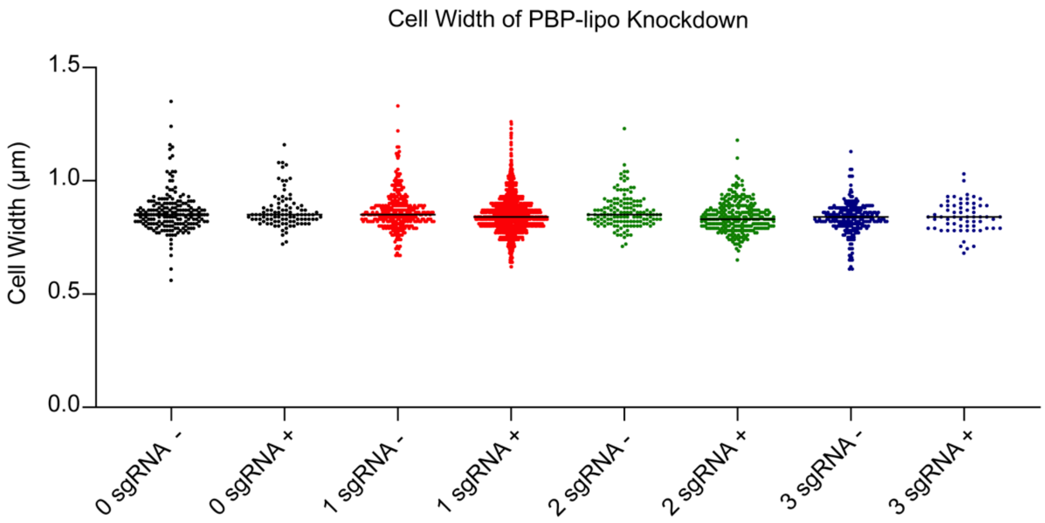


B

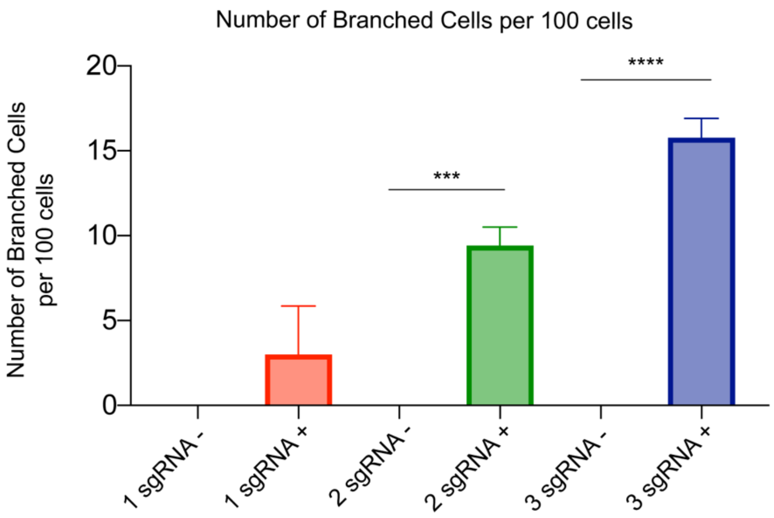


**Supplemental Figure 6. Morphology of PBP-lipo knockdown cells. (A)** Cell width measurements of the 0, 1, 2, and 3 sgRNA PBP-lipo knockdown strains. "-" symbolizes strains grown without ATc. "+" indicates strains grown with ATc. PBP-lipo was induced for knockdown for 18 hr and cultures were visualized on the microscope and analyzed using the GEMATRIA and MOMIA programs (34). **(B)** Number of branched cells per 100 cells visualized under light microscopy. **** $p < 0.001$, *** $p = 0.001$.

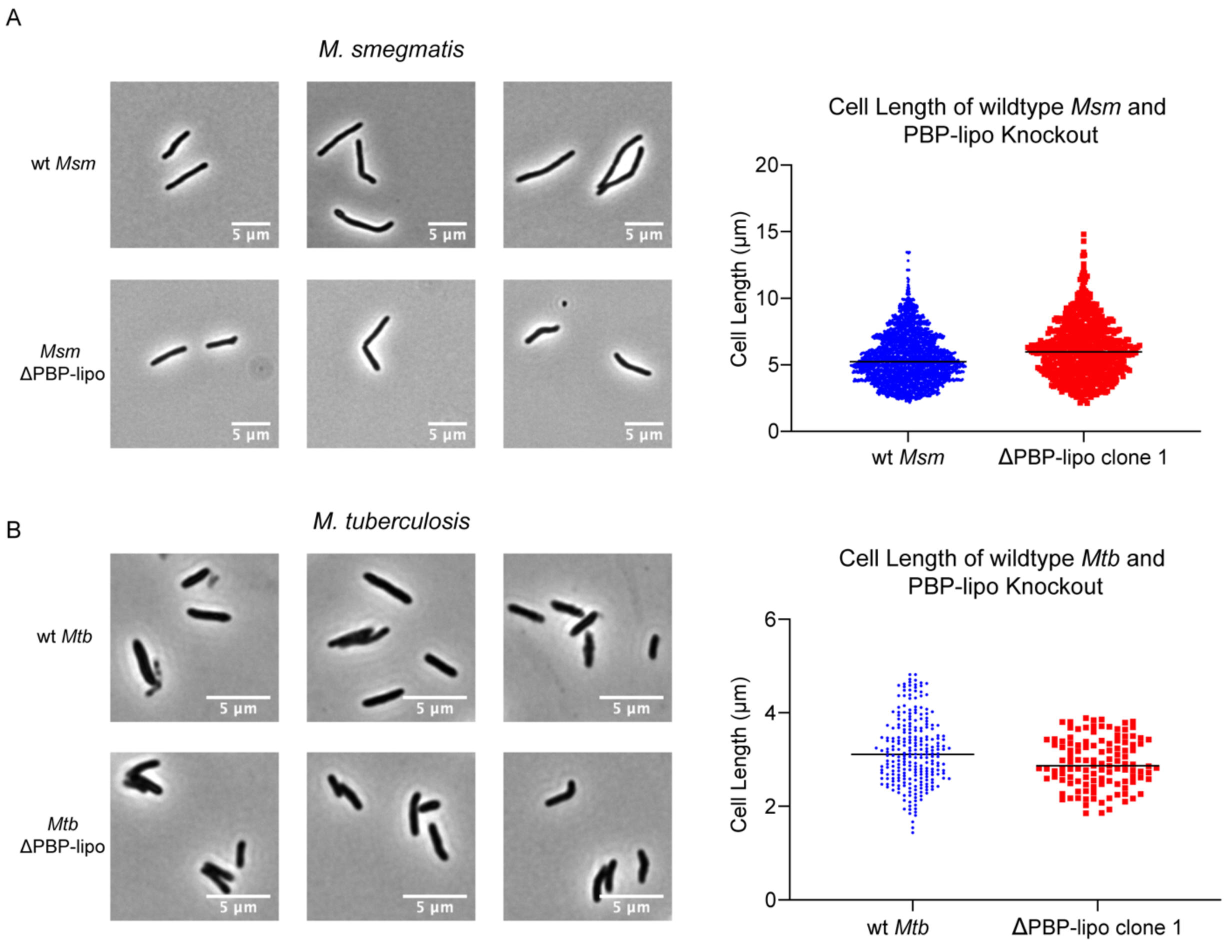


**Supplemental Figure 7. Knockout of PBP-lipo does not alter morphology of *Msm* and *Mtb* cells. (A)** (*left*) Microscopy images of wildtype *Msm* and ΔPBP-lipo strain. (*right*) Quantification of cell length **(B)** (*left*) Microscopy images of wildtype *Mtb* and ΔPBP-lipo mutant. (*right*) Quantification of cell length All morphological measurements were conducted using the GEMATRIA and MOMIA pipelines (34).

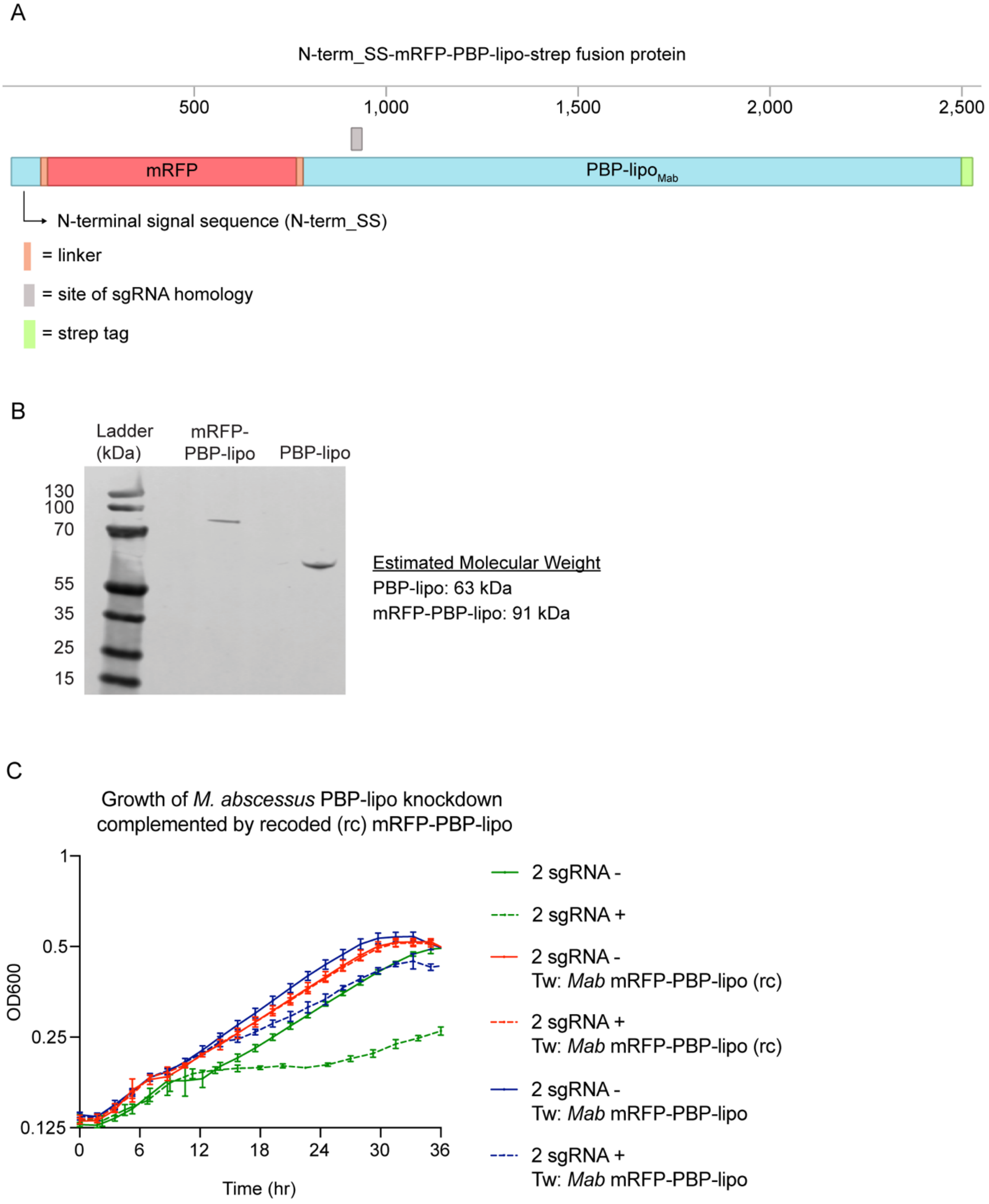


**Supplemental Figure 8. mRFP-PBP-lipo is a functional fusion protein. (A)** Schematic of mRFP-PBP-lipo-strep fusion protein. **(B)** Western Blot using α-strep antibody on strains expressing PBP-lipo-strep and mRFP-PBP-lipo-strep. **(C)** Growth curve showing complementation with mRFP-PBP-lipo fusion proteins.

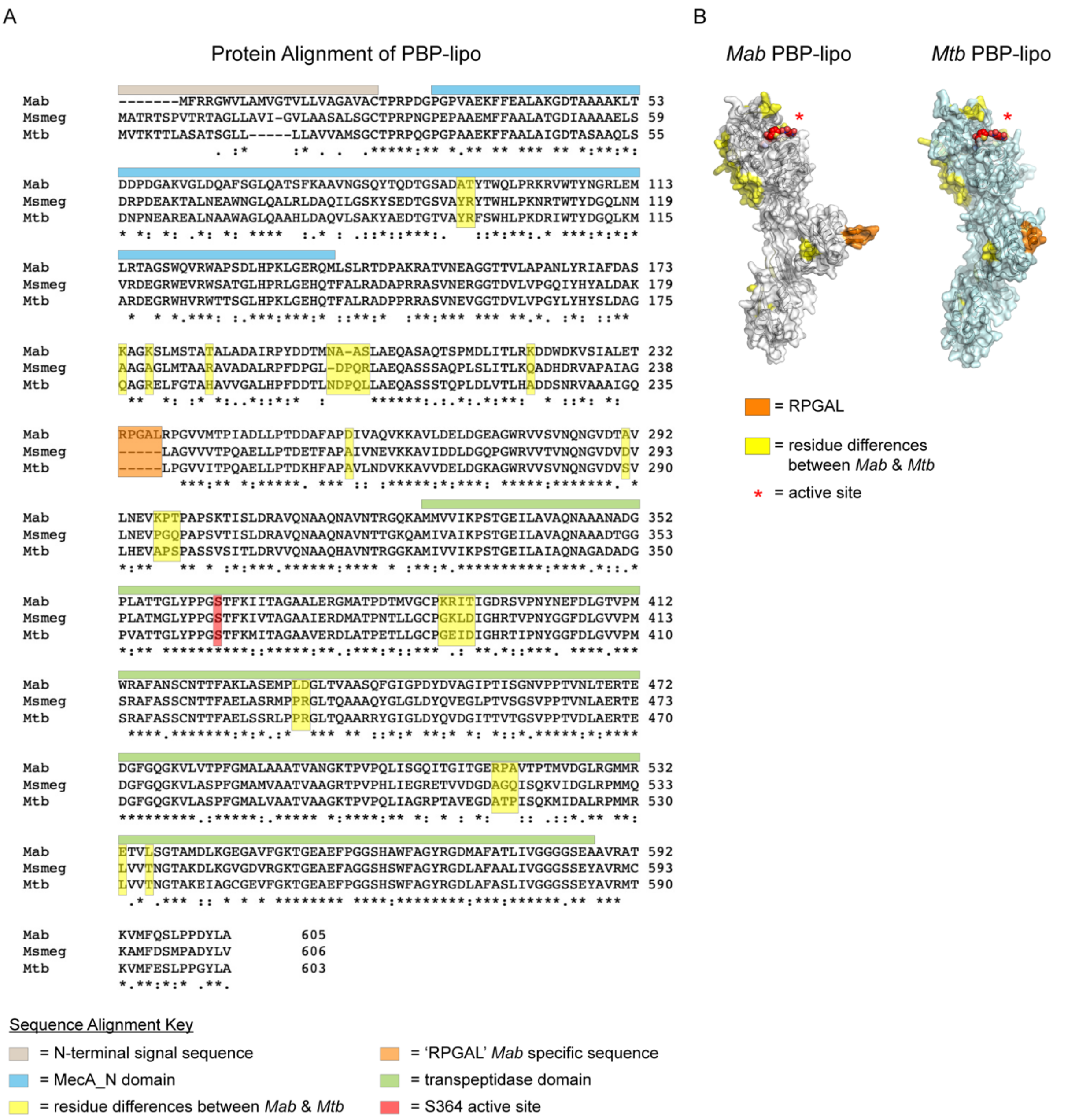


**Supplemental Figure 9. Structural Comparison of PBP-lipo across mycobacteria. (A)** Protein alignment of *Mab, Msm, Mtb, M. bovis,* and *M. leprae* PBP-lipo homologs. Orange box indicates 5 additional amino acids "RGPAL" present in *Mab's* PBP-lipo. Residues highlighted in yellow indicate significant differences in amino acid identity between *Mab* and *Mtb*. **(B)** *in silico* generated model of *Mab* and *Mtb* PBP-lipo structures. Schematized β-lactam is depicted binding the predicted active site of PBP-lipo. Residues in orange highlight the additional "RPGAL" amino acids present in *Mab*. Residues in yellow correspond to amino acid differences between *Mab* and *Mtb* PBP-lipo.

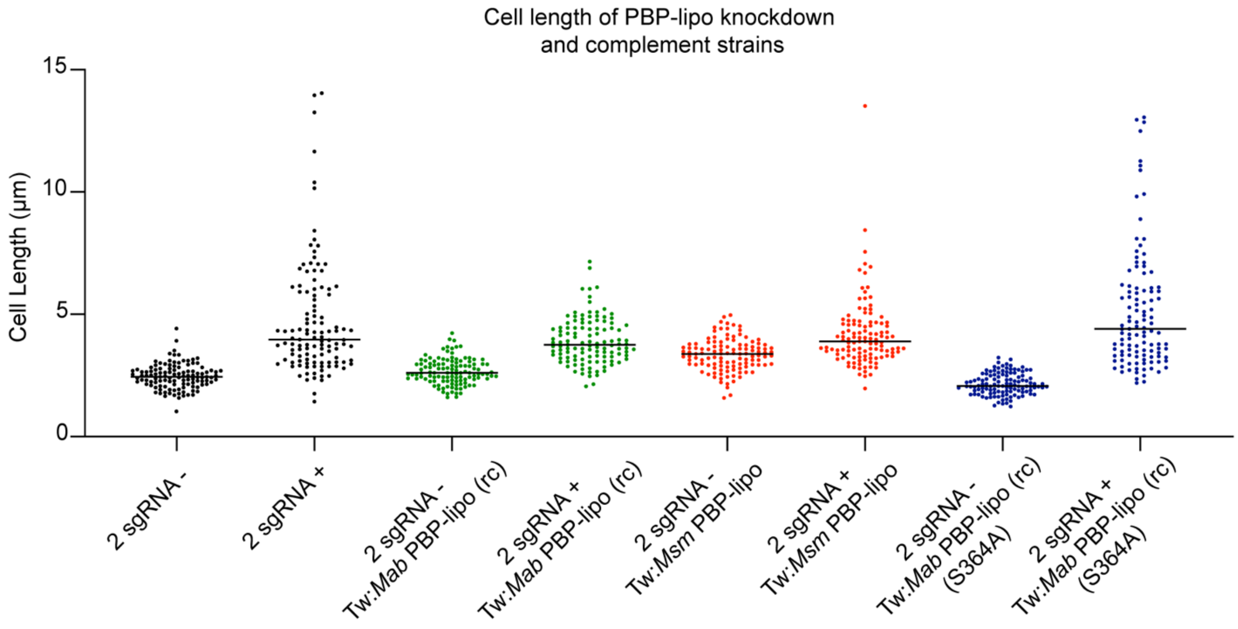


**Supplemental Figure 10. Cell length of PBP-lipo knockdown and complement strains.**

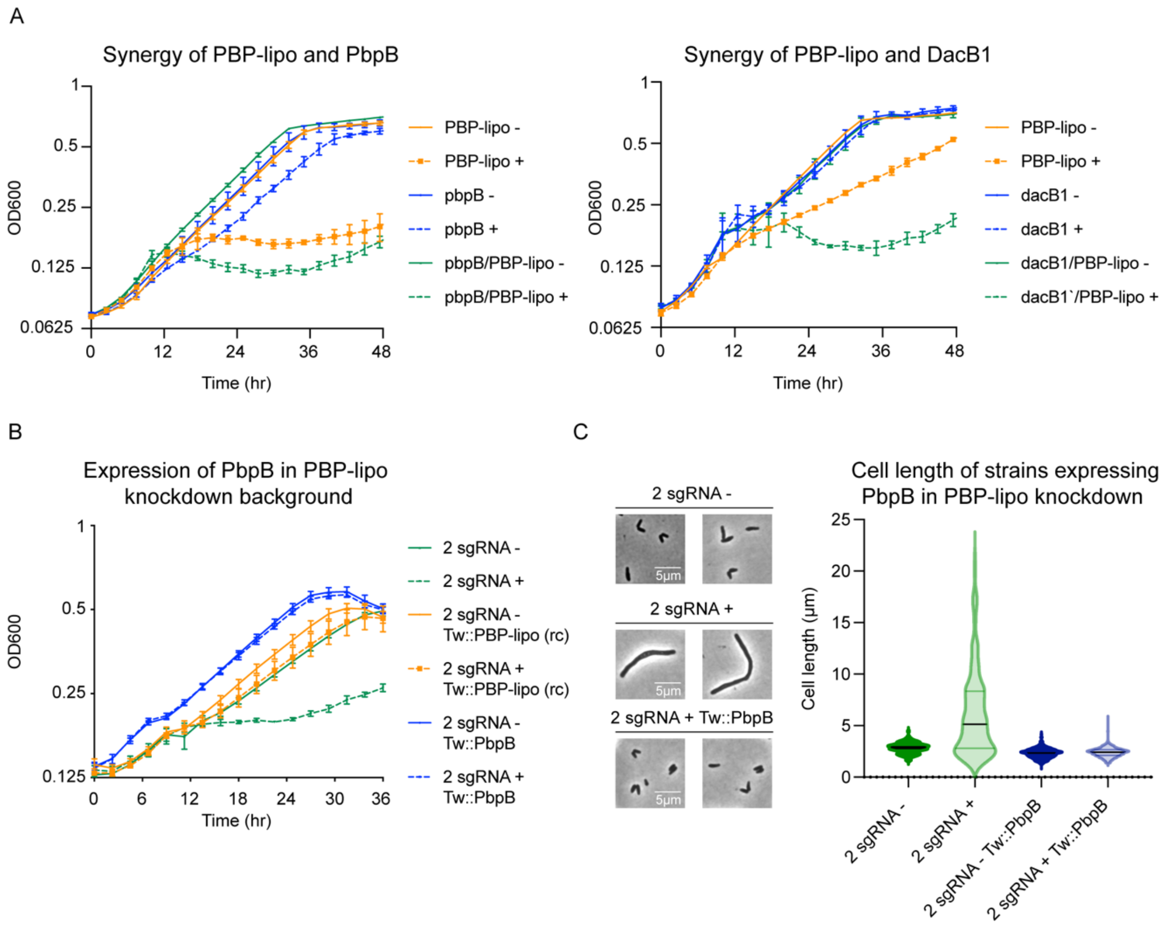


**Supplemental Figure 11. Genetic interactions of PBP-lipo and PBPs. (A)** Genetic synergy of PBP-lipo/PbpB and PBP-lipo/DacB1 in liquid culture. **(B)** Growth curve of PBP-lipo knockdown overexpressing PbpB. **(C)** (left) Microscopy images of 2 sgRNA PBP-lipo knockdown and PbpB complement strains (right) Cell lengths of 2 sgRNA PBP-lipo knockdown cells and PbpB complement strain.

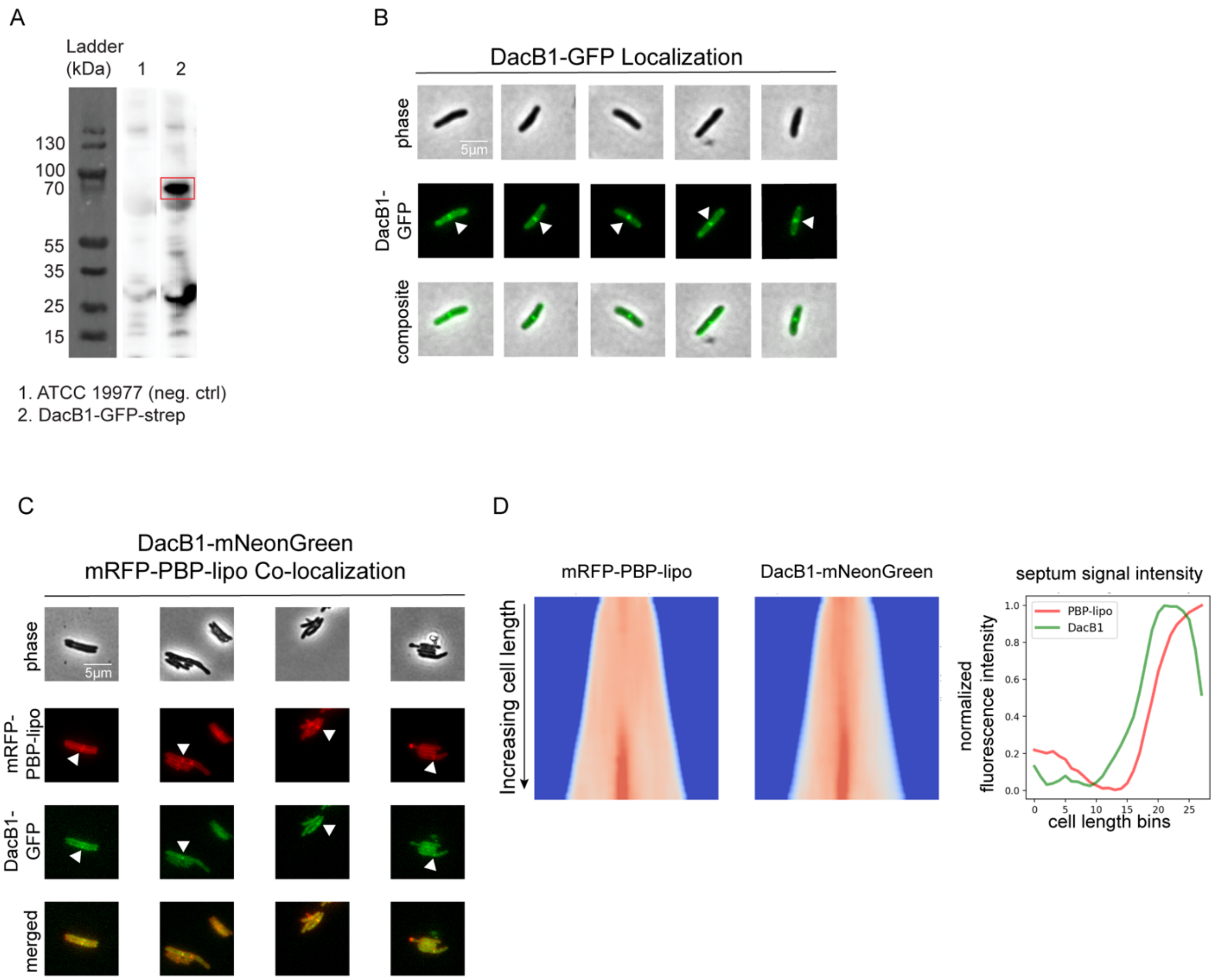


**Supplemental Figure 12: DacB1 localizes to the septum of *Mab* and co-localizes with PBP-lipo. (A)** Western Blot using α-strep antibody on strain expressing DacB1-GFP-strep. Estimated MW of DacB1-GFP is 72 kDa. **(B)** Microscopy images of DacB1 C-terminally tagged with GFP. White arrows point to septal localization of DacB1 **(C)** Microscopy images of N-terminally tagged mRFP-PBP-lipo and C-terminally tagged DacB1-GFP. 'Merged' images show overlay of red and green channels **(D)** *(left)* Demograph of mRFP-PBP-lipo and DacB1-mNeonGreen and *(right)* Fluorescence signal arranged by increasing cell length.

A

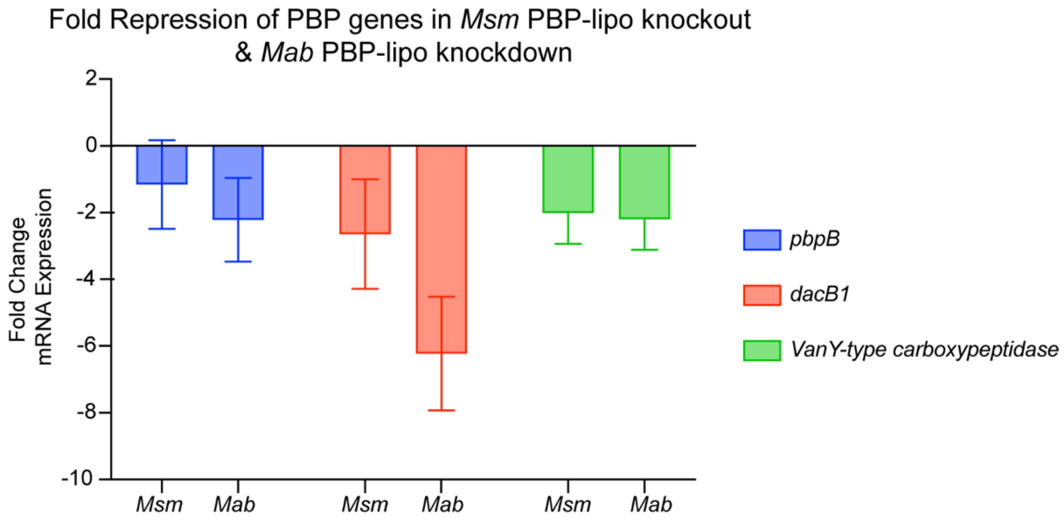


B

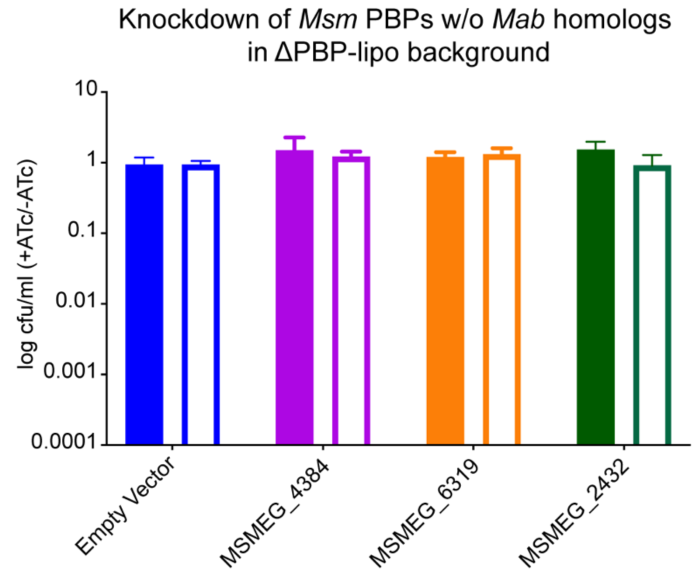


**Supplemental Figure 13. Genetic Interactions of PBPs in *Msm* and *Mab* (A)** Fold repression of genes with genetic synergy in *Msm* knockout and *Mab* knockdown. **(B)** Genetic synergy of *Msm* PBP-lipo with PBPs that lack homologs in *Mab*.

A

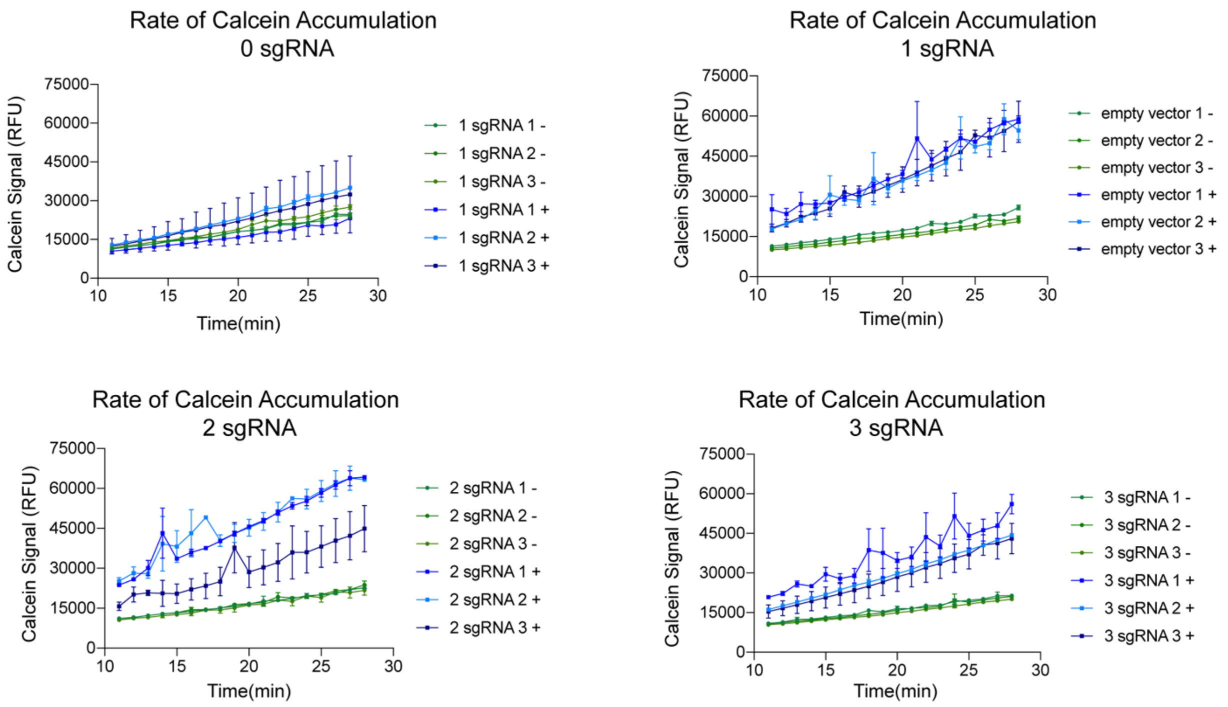


B

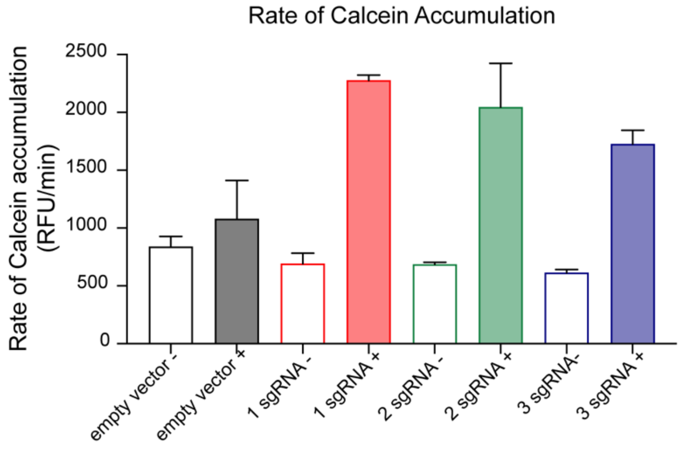


**Supplemental Figure 14. Knockdown of PBP-lipo increases rate of calcein accumulation in *Mab*.** **(A)** Measurement of calcein signal in cultures that were induced for PBP-lipo knockdown with either 1, 2, or 3 sgRNAs. **(B)** Rate of calcein signal accumulation.

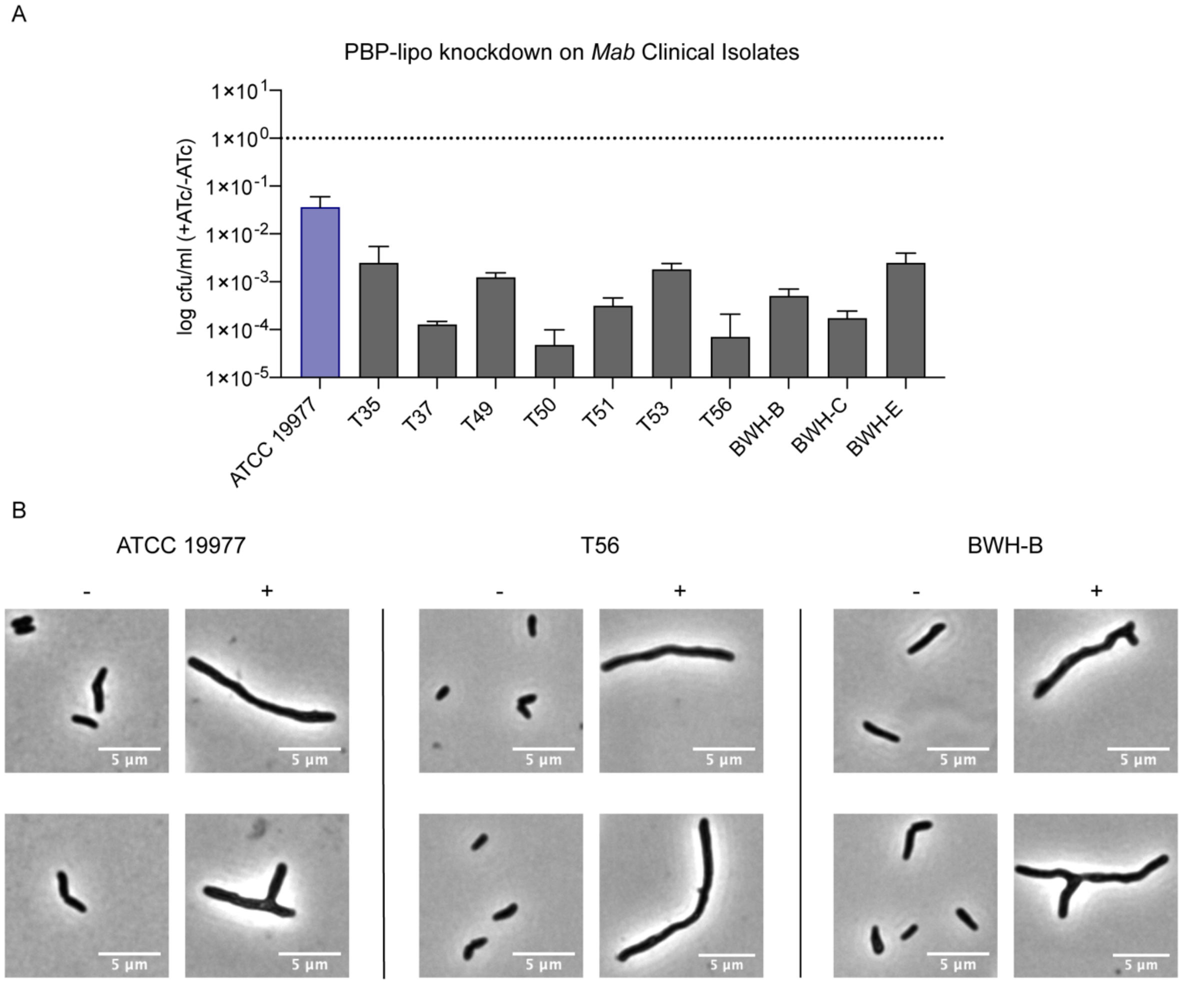


**Supplemental Figure 15. Knockdown of PBP-lipo impairs growth in clinical isolates. (A)** CFU of *Mab* clinical isolate with 3 sgRNA plasmid targeting PBP-lipo. Cultures were spotted on 7H10 plates +/- ATc. **(B)** Microscopy images ATCC 19977, T56, and BWH-B strains induced for PBP-lipo knockdown.

**Supplemental Table 1: Summary of *Mab subsp. abscessus* ATCC 19977 TnSeq Libraries**

| | **Total Counts** | **% Saturation** |
|---|---|---|
| Library #1 | 1176998 | 54.2 |
| Library #2 | 3600892 | 62.3 |
| Library #3 | 4080756 | 64.1 |

**Supplemental Table 3: List of essential genes in *Mab* with *non-essential* orthologs in *Mtb***

| ***Mab* ortholog** | **H37Rv ortholog** | **gene** | **E-value** | **% aa identity** | **annotation of H37Rv gene** |
|---|---|---|---|---|---|
| MAB_2129 | Rv2124c | metH | 1.00E-174 | 34 | 5-methyltetrahydrofolate--homocystein methyltransferase MetH (methionine synthase, vitamin-B12 dependent isozyme) (ms) |
| MAB_3675 | Rv3318 | sdhA | 0 | 90 | Probable succinate dehydrogenase (flavoprotein subunit) SdhA (succinic dehydrogenase) (fumarate reductase) (fumarate dehydrogenase) (fumaric hydrogenase) |
| MAB_3686c | Rv0066c | icd2 | 0 | 81 | Probable isocitrate dehydrogenase [NADP] Icd2 (oxalosuccinate decarboxylase) (IDH) (NADP+-specific ICDH) (IDP) |
| MAB_3167c | Rv2864c | - | 0 | 64 | Possible penicillin-binding lipoprotein |
| MAB_1920 | Rv2222c | glnA2 | 0 | 86 | Probable glutamine synthetase GlnA2 (glutamine synthase) (GS-II) |
| MAB_1662c | Rv2391 | sirA | 0 | 78 | Ferredoxin-dependent sulfite reductase SirA |
| MAB_1563 | Rv2474c | - | 9.00E-76 | 55 | hypothetical protein |
| MAB_1089 | Rv0996 | - | 5.00E-68 | 49 | Probable conserved transmembrane protein |
| MAB_4049c | Rv0489 | gpm1 | 1.00E-137 | 79 | Probable phosphoglycerate mutase 1 Gpm1 (phosphoglyceromutase) (PGAM) (BPG-dependent PGAM) |
| MAB_1484 | Rv1340 | rphA | 7.00E-149 | 85 | Probable ribonuclease RphA (RNase PH) (tRNA nucleotidyltransferase) |
| MAB_3602c | Rv3256c | - | 2.00E-99 | 51 | hypothetical protein |
| MAB_3110 | Rv2788 | sirR | 4.00E-125 | 78 | Probable transcriptional repressor SirR |
| MAB_3673 | Rv3316 | sdhC | 2.00E-58 | 75 | Probable succinate dehydrogenase (cytochrome B-556 subunit) SdhC (succinic dehydrogenase) (fumarate reductase) (fumarate dehydrogenase) (fumaric hydrogenase) |
| MAB_3995 | Rv0505c | serB1 | 8.00E-142 | 82 | Possible phosphoserine phosphatase SerB1 (PSP) (O-phosphoserine phosphohydrolase) (pspase) |
| MAB_2848c | Rv2552c | aroE | 4.00E-94 | 60 | Probable shikimate 5-dehydrogenase AroE (5-dehydroshikimate reductase) |
| MAB_1513 | Rv2523c | acpS | 5.00E-68 | 74 | holo-[acyl-carrier protein] synthase AcpS (holo-ACP synthase) (CoA:APO-[ACP]pantetheinephosphotransferase) (CoA:APO-[acyl-carrier protein]pantetheinephosphotransferase) |
| MAB_3676 | Rv3319 | sdhB | 8.00E-167 | 84 | Probable succinate dehydrogenase (iron-sulphur protein subunit) SdhB (succinic dehydrogenase) (fumarate reductase) (fumarate dehydrogenase) (fumaric hydrogenase) |
| MAB_4954c | Rv3923c | rnpA | 3.00E-27 | 51 | Ribonuclease P protein component RnpA (RNaseP protein) (RNase P protein) (protein C5) |
| MAB_1446 | Rv1303 | - | 3.00E-41 | 57 | hypothetical protein |
| MAB_4471 | Rv0236A | - | 3.00E-23 | 68 | Small secreted protein |

**Supplemental Table 4: TnSeq Summary of *Msm* Libraries**

| Strain | Total counts | Saturation | # of conditional essentials compared to wt |
|---|---|---|---|
| mc$^2$155 -1 | 1805936 | 0.519 | - |
| mc$^2$155 -2 | 1264512 | 0.534 | - |
| ΔPBP-lipo-1 | 1389005 | 0.628 | 0 |

**Supplemental Table 5: MIC (µg/ml) of wildtype mc$^{2}$155 & ΔPBP-lipo**

| Antibiotic | -ATc | +ATc | Fold Difference |
|---|---|---|---|
| **Cell Wall** | | | |
| Ampicillin | 8 | 16 | 0.5 |
| Amoxicillin | 4 | 4 | 1 |
| Faropenem | >64 | >64 | N/A |
| Cefoxitin | 32 | 32 | 1 |
| Vancomycin | 4 | 2 | 2 |
| Meropenem | >64 | >64 | N/A |
| Imipenem | >64 | >64 | “ “ |
| Ticarcillin | >64 | >64 | “ “ |
| Ceftazidime | >64 | >64 | “ “ |
| Cephalexin | >64 | >64 | “ “ |
| Ceftazidime | >64 | >64 | N/A |
| D-Cycloserine | 32 | 32 | 1 |
| Ethambutol | 0.25 | 0.25 | “ “ |
| Isoniazid | 16 | 16 | “ “ |
| **Ribosome** | | | |
| Clarithromycin | 0.25 | <0.125 | N/A |
| Erythromycin | 4 | 1 | 4 |
| Amikacin | <0.125 | 0.125 | N/A |
| Clindamycin | >64 | 64 | “ “ |
| **RNA Polymerase** | | | |
| Rifampicin | 16 | 8 | 2 |
| **DNA Gyrase** | | | |
| Ofloxacin | 0.25 | 0.25 | 1 |
| **Other** | | | |
| Pyrazinamide | >64 | >64 | N/A |
| Pretomanid | >64 | >64 | “ “ |

**Supplemental Table 6: MIC (μg/ml) of wildtype H37Rv & ΔPBP-lipo**

| | MIC | | |
|---|---|---|---|
| **Antibiotic** | **wt H37Rv** | **ΔPBP-lipo** | **Fold Difference** |
| **Cell Wall** | | | |
| Ampicillin | 8 | 16 | 0.5 |
| Amoxicillin | 4 | 4 | 1 |
| Faropenem | >64 | >64 | N/A |
| Cefoxitin | 32 | 32 | 1 |
| **Ribosome** | | | |
| Clarithromycin | 0.25 | <0.125 | N/A |
| Erythromycin | 4 | 1 | 4 |
| Amikacin | <0.125 | 0.125 | N/A |
| **RNA Polymerase** | | | |
| Rifampicin | 16 | 8 | 2 |
| **DNA Gyrase** | | | |
| Ofloxacin | 0.25 | 0.25 | 1 |
