## Supplemental Table 7 for "High-density transposon mutagenesis in *Mycobacterium abscessus* identifies an essential penicillin-binding lipo-protein (PBP-lipo) involved in septal peptidoglycan synthesis and antibiotic sensitivity"

**Supplemental Table 7: Bacterial Strains Used**

| **Strain #** | **Strain Base** | **Genotype** | **Additional Information** |
| --- | --- | --- | --- |
| CA176 | *Mab* | Mycobacterium abscessus subsp. abscessus ATCC 19977 | wildtype *Mab* reference strain |
| CA401 | *Mab* | ATCC 19977 L5::CT296 pUV15-Tet-dCas9-empty sgRNA | CRISPRi plasmid carrying empty sgRNA used for control experiments |
| CA409 | *Mab* | ATCC 19977 L5::cr23 pUV15-Tet-dCas9-cr23-*(rpoB_Mab_)* | CRISPRi plasmid carrying 1 sgRNA targeting *rpoB* |
| CA415 | *Mab* | ATCC 19977 L5::cr41 pUV15-Tet-dCas9-cr41-*(gyrB_Mab_)* | CRISPRi plasmid carrying 1 sgRNA targeting *gyrB* |
| CA417 | *Mab* | ATCC 19977 L5::cr43 pUV15-Tet-dCas9-cr43-*(secY_Mab_)* | CRISPRi plasmid carrying 1 sgRNA targeting *secY* |
| CA522 | *Mab* | ATCC 19977 L5::cr36 pUV15-Tet-dCas9-cr36-*(glnA2_Mab_)* | CRISPRi plasmid carrying 1 sgRNA targeting *glnA2* |
| CA523 | *Mab* | ATCC 19977 L5::cr61 pUV15-Tet-dCas9-cr61-*(icd2_Mab_)* | CRISPRi plasmid carrying 1 sgRNA targeting *icd2* |
| CA524 | *Mab* | ATCC 19977 L5::cr63 pUV15-Tet-dCas9-cr63-*(sdhA_Mab_)* | CRISPRi plasmid carrying 1 sgRNA targeting *sdhA* |
| CA525 | *Mab* | ATCC 19977 L5::cr40 pUV15-Tet-dCas9-cr40-*(MAB_4471)* | CRISPRi plasmid carrying 1 sgRNA targeting *MAB_4471* |
| CA410 | *Mab* | ATCC 19977 L5::cr25 pUV15-Tet-dCas9-cr25-*(PBP-lipo_Mab_)* | CRISPRi plasmid carrying 1 sgRNA targeting PBP-lipo |
| CA453 | *Mab* | ATCC 19977 L5::pCRC1 pUV15-Tet-dCas9-cr25+cr26 *(PBP-lipo_Mab_)* | CRISPRi plasmid carrying 2 sgRNAs targeting PBP-lipo |
| CA450 | *Mab* | ATCC 19977 L5::pCRC2 pUV15-Tet-dCas9-cr25+cr26+cr56 *(PBP-lipo_Mab_)* | CRISPRi plasmid carrying 3 sgRNAs targeting PBP-lipo |
| CA526 | *Mab* | ATCC 19977 L5::pCA276.2 pUV15-N-signal_sequence-mRFP-PBP-lipo_Mab_-strep | N-terminal tag of PBP-lipo w/ mRFP for visualization |
| CA553 | *Mab* | ATCC 19977 L5::308 pUV15-dacB1*_Mab_*-GFPmut3 | C-terminal tag of PBP-lipo w/ GFPmut3 for visualization |
| CA519 | *Mab* | ATCC 19977 Tw::pCA272 MOP-PBP-lipo_Mab_-strep | Overexpression of PBP-lipo-strep |
| CA467 | *Mab* CA453 | ATCC 19977 L5::pCRC1 pUV15-Tet-dCas9-cr25+cr26 *(PBP-lipo_Mab_)* Tw::pCA242 MOP-PBP_lipo_Msm_-myc | CRISPRi plasmid carrying 2 sgRNAs targeting *PBP-lipo* + Plasmid that constitutively expresses PBP-lipo_Msm_ |
| CA435 | *Mab* | ATCC 19977 L5:pCRC13 pUV15-Tet-dCas9-cr25 *(PBP-lipo_Mab_)* + empty guide | CRISPRi plasmid carrying sgRNAs targeting *PBP-lipo* and empty guide |
| CA527 | *Mab* | ATCC 19977 L5:cr67 pUV15-Tet-dCas9-cr67-*(ponA1_Mab_)* | CRISPRi plasmid carrying 1 sgRNAs targeting *ponA1* |
| CA535 | *Mab* | ATCC 19977 L5:pCRC5 pUV15-Tet-dCas9-cr67-*(ponA1_Mab_)* + cr25 *(PBP-lipo_Mab_)* | CRISPRi plasmid carrying sgRNAs targeting PBP-lipo and ponA1 |
| CA528 | *Mab* | ATCC 19977 L5:cr68 pUV15-Tet-dCas9-cr68 *(ponA2_Mab_)* | CRISPRi plasmid carrying 1 sgRNAs targeting *ponA2* |
| CA536 | *Mab* | ATCC 19977 L5:pCRC6 L5:cr68 pUV15-Tet-dCas9-cr68 *(ponA2_Mab_)* + cr25 *(PBP-lipo_Mab_)* | CRISPRi plasmid carrying sgRNAs targeting PBP-lipo and *ponA2* |
| CA529 | *Mab* | ATCC 19977 L5:cr69 pUV15-Tet-dCas9-cr69-*(pbpA_Mab_)* | CRISPRi plasmid carrying 1 sgRNAs targeting *pbpA* |
| CA537 | *Mab* | ATCC 19977 L5:pCRC7 pUV15-Tet-dCas9-cr69-*(pbpA_Mab_)* + cr25 *(PBP-lipo_Mab_)* | CRISPRi plasmid carrying sgRNAs targeting *PBP-lipo* and *pbpA* |
| CA530 | *Mab* | ATCC 19977 L5:cr70 pUV15-Tet-dCas9-cr70-*(pbpB_Mab_)* | CRISPRi plasmid carrying 1 sgRNAs targeting *pbpB* |
| CA538 | *Mab* | ATCC 19977 L5:pCRC8 pUV15-Tet-dCas9-cr70-*(pbpB_Mab_)* + cr25 *(PBP-lipo_Mab_)* | CRISPRi plasmid carrying sgRNAs targeting *PBP-lipo* and *pbpB* |
| CA531 | *Mab* | ATCC 19977 L5:cr71 pUV15-Tet-dCas9-cr71-*(dacB1_Mab_)* | CRISPRi plasmid carrying 1 sgRNAs targeting *dacB1* |
| CA434 | *Mab* | ATCC 19977 L5:pCRC9 pUV15-Tet-dCas9-cr71-*(dacB1_Mab_)* + cr25 *(PBP-lipo_Mab_)* | CRISPRi plasmid carrying sgRNAs targeting *PBP-lipo* and *dacB1* |
| CA532 | *Mab* | ATCC 19977 L5:cr72 pUV15-Tet-dCas9-cr72-*(dacB2_Mab_)* | CRISPRi plasmid carrying 1 sgRNAs targeting *dacB2* |
| CA540 | *Mab* | ATCC 19977 L5:pCRC10 pUV15-Tet-dCas9-cr72-*(dacB2_Mab_)* + cr25 *(PBP-lipo_Mab_)* | CRISPRi plasmid carrying sgRNAs targeting *PBP-lipo* and *dacB2* |
| CA533 | *Mab* | ATCC 19977 L5:cr73 pUV15-Tet-dCas9-cr73-*(carboxypeptidase_Mab_)* | CRISPRi plasmid carrying 1 sgRNAs targeting *carboxypeptidase* |
| CA541 | *Mab* | ATCC 19977 L5:pCRC11 pUV15-Tet-dCas9-cr73-*(carboxypeptidase_Mab_)* + cr25 *(PBP-lipo_Mab_)* | CRISPRi plasmid carrying sgRNAs targeting *PBP-lipo* and *carboxypeptidase* |
| CA534 | *Mab* | ATCC 19977 L5:cr74 pUV15-Tet-dCas9-cr74-*(VanY-type carboxypeptidase_Mab_)* | CRISPRi plasmid carrying 1 sgRNAs targeting *VanY-type carboxypeptidase* |
| CA542 | *Mab* | ATCC 19977 L5:pCRC12 pUV15-Tet-dCas9-cr74-*(VanY-type carboxypeptidase_Mab_)* + cr25 *(PBP-lipo_Mab_)* | CRISPRi plasmid carrying sgRNAs targeting *PBP-lipo* and *VanY-type carboxypeptidase* |
| CA543 | *Mab* | T35 L5::pCRC2 pUV15-Tet-dCas9-cr25+cr26+cr56 *(PBP-lipo_Mab_)* | *Mab* clinical isolate with CRISPRi plasmid with 3 sgRNAs targeting PBP-lipo |
| CA544 | *Mab* | T37 L5::pCRC2 pUV15-Tet-dCas9-cr25+cr26+cr56 *(PBP-lipo_Mab_)* | *Mab* clinical isolate with CRISPRi plasmid with 3 sgRNAs targeting PBP-lipo |
| CA545 | *Mab* | T49 L5::pCRC2 pUV15-Tet-dCas9-cr25+cr26+cr56 *(PBP-lipo_Mab_)* | *Mab* clinical isolate with CRISPRi plasmid with 3 sgRNAs targeting PBP-lipo |
| CA546 | *Mab* | T50 L5::pCRC2 pUV15-Tet-dCas9-cr25+cr26+cr56 *(PBP-lipo)* | *Mab* clinical isolate with CRISPRi plasmid with 3 sgRNAs targeting PBP-lipo |
| CA547 | *Mab* | T51 L5::pCRC2 pUV15-Tet-dCas9-cr25+cr26+cr56 *(PBP-lipo)* | *Mab* clinical isolate with CRISPRi plasmid with 3 sgRNAs targeting PBP-lipo |
| CA548 | *Mab* | T53 L5::pCRC2 pUV15-Tet-dCas9-cr25+cr26+cr56 *(PBP-lipo_Mab_)* | *Mab* clinical isolate with CRISPRi plasmid with 3 sgRNAs targeting PBP-lipo |
| CA549 | *Mab* | T56 L5::pCRC2 pUV15-Tet-dCas9-cr25+cr26+cr56 *(PBP-lipo_Mab_)* | *Mab* clinical isolate with CRISPRi plasmid with 3 sgRNAs targeting PBP-lipo |
| CA550 | *Mab* | BWH-B L5::pCRC2 pUV15-Tet-dCas9-cr25+cr26+cr56 *(PBP-lipo_Mab_)* | *Mab* clinical isolate with CRISPRi plasmid with 3 sgRNAs targeting PBP-lipo |
| CA551 | *Mab* | BWH-C L5::pCRC2 pUV15-Tet-dCas9-cr25+cr26+cr56 *(PBP-lipo_Mab_)* | *Mab* clinical isolate with CRISPRi plasmid with 3 sgRNAs targeting PBP-lipo |
| CA552 | *Mab* | BWH-E L5::pCRC2 pUV15-Tet-dCas9-cr25+cr26+cr56 *(PBP-lipo_Mab_)* | *Mab* clinical isolate with CRISPRi plasmid with 3 sgRNAs targeting PBP-lipo |
| IDW9 | *Mtb* | wildtype H37Rv | wildtype *Mtb* |
| IDW7 | *Mtb* | H37Rv PBP-lipo_Mtb_::zeoR (clone B) | PBP-lipo knockout in *Mtb* |
| IDW8 | *Mtb* | H37Rv PBP-lipo_Mtb_::zeoR (clone D) |  |
| CA430 | *Msm* | wildtype mc2 155 | wildtype *M. smegmatis* |
| CA464 | *Msm* | mc2 155 PBP-lipo_Msm_::zeoR (clone 1) | PBP-lipo knockout in *Mtb* |
| CA464 | *Msm* | mc2 155 PBP-lipo_Msm_::zeoR (clone 2) | PBP-lipo knockout in *Mtb* |
| CA480 | *Msm* | mc2 155 L5::pcr86 pUV15-Tet-dCas9-cr86-*(ponA1_Msm_)* | CRISPRi plasmid carrying 1 sgRNA targeting *ponA1* (strong guide) |
| CA481 | *Msm* | mc2 155 L5::pcr87 pUV15-Tet-dCas9-cr87-*(ponA1 _Msm_)* | CRISPRi plasmid carrying 1 sgRNA targeting *ponA1* (weak guide) |
| CA482 | *Msm* | mc2 155 L5::pcr88 pUV15-Tet-dCas9-cr88-*(ponA2 _Msm_)* | CRISPRi plasmid carrying 1 sgRNA targeting *ponA2* |
| CA483 | *Msm* | mc2 155 L5::pcr89 pUV15-Tet-dCas9-cr89-*(MSMEG_4384)* | CRISPRi plasmid carrying 1 sgRNA targeting *MSMEG_4384* |
| CA484 | *Msm* | mc2 155 L5::pcr90 pUV15-Tet-dCas9-cr90-*(pbpA _Msm_)* | CRISPRi plasmid carrying 1 sgRNA targeting *pbpA* |
| CA485 | *Msm* | mc2 155 L5::pcr91 pUV15-Tet-dCas9-cr91-*(pbpB _Msm_)* | CRISPRi plasmid carrying 1 sgRNA targeting *pbpB* |
| CA486 | *Msm* | mc2 155 L5::pcr92 pUV15-Tet-dCas9-cr92-*(PBP-lipo _Msm_)* | CRISPRi plasmid carrying 1 sgRNA targeting *PBP-lipo* |
| CA487 | *Msm* | mc2 155 L5::pcr93 pUV15-Tet-dCas9-cr93-*(MSMEG_6319)* | CRISPRi plasmid carrying 1 sgRNA targeting *MSMEG_6319* |
| CA488 | *Msm* | mc2 155 L5::pcr94 pUV15-Tet-dCas9-cr94-*(dacB1 _Msm_)* | CRISPRi plasmid carrying 1 sgRNA targeting *dacB1* |
| CA489 | *Msm* | mc2 155 L5::pcr95 pUV15-Tet-dCas9-cr95-*(dacB2 _Msm_)* | CRISPRi plasmid carrying 1 sgRNA targeting *dacB2* |
| CA490 | *Msm* | mc2 155 L5::pcr96 pUV15-Tet-dCas9-cr96-*(MSMEG_2432)* | CRISPRi plasmid carrying 1 sgRNA targeting *MSMEG_2432* |
| CA491 | *Msm* | mc2 155 L5::pcr97 pUV15-Tet-dCas9-cr97-*(carboxypeptidase_Msm_)* | CRISPRi plasmid carrying 1 sgRNA targeting *carboxypeptidase* |
| CA492 | *Msm* | mc2 155 L5::pcr98 pUV15-Tet-dCas9-cr98-*(VanY-carboxypeptidase_Msm_)* | CRISPRi plasmid carrying 1 sgRNA targeting *VanY-type carboxypeptidase* |
| CA493 | *Msm* | mc2 155 L5::pCT295 pUV15-Tet-dCas9-cr23-*(empty guide)* | CRISPRi plasmid carrying empty guide |
| CA494 | *Msm*  CA464 | mc2 155 PBP-lipo_Msm_::zeoR L5::pcr86 pUV15-Tet-dCas9-cr86-*(ponA1_Msm_)* | CRISPRi plasmid carrying 1 sgRNA targeting *ponA1* (strong guide) |
| CA495 | *Msm*  CA464 | mc2 155 PBP-lipo_Msm_::zeoR L5::pcr87 pUV15-Tet-dCas9-cr87-*(ponA1_Msm_)* | CRISPRi plasmid carrying 1 sgRNA targeting *ponA1* (weak guide) |
| CA496 | *Msm*  CA464 | mc2 155 PBP-lipo_Msm_::zeoR L5::pcr88 pUV15-Tet-dCas9-cr88-*(ponA2 _Msm_)* | CRISPRi plasmid carrying 1 sgRNA targeting *ponA2* |
| CA497 | *Msm*  CA464 | mc2 155 PBP-lipo_Msm_::zeoR L5::pcr89 pUV15-Tet-dCas9-cr89-*(MSMEG_4384)* | CRISPRi plasmid carrying 1 sgRNA targeting *MSMEG_4384* |
| CA498 | *Msm*  CA464 | mc2 155 PBP-lipo_Msm_::zeoR L5::pcr90 pUV15-Tet-dCas9-cr90-*(pbpA_Msm_)* | CRISPRi plasmid carrying 1 sgRNA targeting *pbpA* |
| CA499 | *Msm*  CA464 | mc2 155 PBP-lipo_Msm_::zeoR L5::pcr91 pUV15-Tet-dCas9-cr91-*(pbpB_Msm_)* | CRISPRi plasmid carrying 1 sgRNA targeting *pbpB* |
| CA500 | *Msm*  CA464 | mc2 155 PBP-lipo_Msm_::zeoR L5::pcr92 pUV15-Tet-dCas9-cr92-*(PBP-lipo_Msm_)* | CRISPRi plasmid carrying 1 sgRNA targeting *PBP-lipo* |
| CA501 | *Msm*  CA464 | mc2 155 PBP-lipo_Msm_::zeoR L5::pcr93 pUV15-Tet-dCas9-cr93-*(MSMEG_6319)* | CRISPRi plasmid carrying 1 sgRNA targeting *MAB_6319* |
| CA502 | *Msm*  CA464 | mc2 155 PBP-lipo_Msm_::zeoR L5::pcr94 pUV15-Tet-dCas9-cr94-*(dacB1_Msm_)* | CRISPRi plasmid carrying 1 sgRNA targeting *dacB1* |
| CA503 | *Msm*  CA464 | mc2 155 PBP-lipo_Msm_::zeoR L5::pcr95 pUV15-Tet-dCas9-cr95-*(dacB2_Msm_)* | CRISPRi plasmid carrying 1 sgRNA targeting *dacB2* |
| CA504 | *Msm*  CA464 | mc2 155 PBP-lipo_Msm_::zeoR L5::pcr96 pUV15-Tet-dCas9-cr96-*(MSMEG_2432)* | CRISPRi plasmid carrying 1 sgRNA targeting *MSMEG_2432* |
| CA505 | *Msm*  CA464 | mc2 155 PBP-lipo_Msm_::zeoR L5::pcr97 pUV15-Tet-dCas9-cr97-*(carboxypeptidase_Msm_)* | CRISPRi plasmid carrying 1 sgRNA targeting *carboxypeptidase* |
| CA506 | *Msm*  CA464 | mc2 155 PBP-lipo_Msm_::zeoR L5::pcr98 pUV15-Tet-dCas9-cr98-*(VanY-carboxypeptidase_Msm_)* | CRISPRi plasmid carrying 1 sgRNA targeting *VanY-type carboxypeptidase* |
| CA507 | *Msm*  CA464 | mc2 155 PBP-lipo_Msm_::zeoR L5::pCT295 pUV15-Tet-dCas9-*(empty guide)* | CRISPRi plasmid carrying empty guide |
| CA563 | *Mab*  CA453 | ATCC 19977 L5::pCRC1 pUV15-Tet-dCas9-cr25+cr26 *(PBP-lipo_Mab_)* Tw::pCA323 MOP-PBP_lipo_Mab_(recode)-strep | CRISPRi plasmid carrying 2 sgRNAs targeting *PBP-lipo* + Plasmid that constitutively expresses recorded *Mab* PBP-lipo |
| CA569 | *Mab*  CA453 | ATCC 19977 L5::pCRC1 pUV15-Tet-dCas9-cr25+cr26 *(PBP-lipo_Mab_)* Tw::pCA324 MOP-PBP_lipo_Mab_(recode)(S364A)-strep | CRISPRi plasmid carrying 2 sgRNAs targeting *PBP-lipo* + Plasmid that constitutively expresses recorded, catalytically dead *Mab* PBP-lipo |
| CA582 | *Mab* | ATCC 19977 L5::pCA276.2 pUV15-N-signal_sequence-mRFP-PBP-lipo_Mab_-strep Tw::pCA341 piMyc-ftsZ-mNeonGreen | Strain used to co-express mRFP-PBP-lipo and ftsZ-mNeonGreen |
| CA583 | *Mab* | ATCC 19977 L5::pCA276.2 pUV15-N-signal_sequence-mRFP-PBP-lipo_Mab_-strep Tw::pCA341 piMyc-ftsZ-mNeonGreen | Strain used to co-express mRFP-PBP-lipo and ftsZ-mNeonGreen |
| CA584 | *Mab*  CA453 | ATCC 19977 L5::pCRC1 pUV15-Tet-dCas9-cr25+cr26 *(PBP-lipo_Mab_)* Tw::pCA353 pUV15-N-signal_sequence-mRFP-PBP-lipo_Mab_(recoded)-strep | Strain used to knockdown native PBP-lipo and express recoded mRFP-PBP-lipo-strep fusion protein |
| CA586 | *Mab*  CA453 | ATCC 19977 L5::pCRC1 pUV15-Tet-dCas9-cr25+cr26 *(PBP-lipo_Mab_)* Tw::pCA353 Tw::pCA353 pUV15-N-signal_sequence-mRFP-PBP-lipo_Mab_-strep | Strain used to knockdown native PBP-lipo and express mRFP-PBP-lipo-strep fusion protein |
| CA588 | *Mab*  CA453 | ATCC 19977 L5::pCRC1 pUV15-Tet-dCas9-cr25+cr26 *(PBP-lipo_Mab_)* Tw::pCA344 MOP-PbpB-strep | Strain used to knockdown native PBP-lipo and express PbpB |
| CA590 | *Mab* | ATCC 19977 L5::pCA276.2 pUV15-N-signal_sequence-mRFP-PBP-lipo_Mab_-strep Tw::pCA342 piMyc-dacB1-mNeonGreen | Strain used to co-express mRFP-PBP-lipo and dacB1-mNeonGreen |
| CA591 | *Mab* | ATCC 19977 L5::pCA276.2 pUV15-N-signal_sequence-mRFP-PBP-lipo_Mab_-strep Tw::pCA342 piMyc-dacB1-mNeonGreen | Strain used to co-express mRFP-PBP-lipo and dacB1-mNeonGreen |
| CA592 | *Mab*  CA453 | ATCC 19977 L5::pCRC1 pUV15-Tet-dCas9-cr25+cr26 *(PBP-lipo_Mab_)* Tw::pCA269-1 natP-GFPmut3-ftsZ | Strain used to knockdown native PBP-lipo and express GFPmut3-ftsZ fusion protein from a natural promoter |
| CA593 | *Mab*  CA453 | ATCC 19977 L5::pCRC1 pUV15-Tet-dCas9-cr25+cr26 *(PBP-lipo_Mab_)* Tw::pCA269-4 natP-GFPmut3-ftsZ | Strain used to knockdown native PBP-lipo and express GFPmut3-ftsZ fusion protein from a natural promoter |
