## Supplemental Table 8 for "High-density transposon mutagenesis in *Mycobacterium abscessus* identifies an essential penicillin-binding lipo-protein (PBP-lipo) involved in septal peptidoglycan synthesis and antibiotic sensitivity"

**Supplemental Table 8: Plasmids Used**

| **Plasmid Number** | **Contents** | **Antibiotic Selection** | **Parental Vector** | **Notes** |
| --- | --- | --- | --- | --- |
| pCT295 | pUV15-Tet-dCas9-*(empty sgRNA)* | kan | CT295 | Gift from Jeremy Rock |
| pCT296 | pUV15-Tet-dCas9-*(empty sgRNA)* | kan | CT296 | Gift from Jeremy Rock­ |
| pCA242 | MOP-PBP-lipo_Msm_-myc | zeo | pCA239 |  |
| pCA243 | MOP-mScarlet-myc | zeo |  |  |
| pCA269-1 | natP-GFPmut3-ftsZ | zeo | pCA243 |  |
| pCA269-4 | natP-GFPmut3-ftsZ | zeo | pCA243 |  |
| pCA272 | MOP-PBP-lipo_Mab_-strep | zeo | pCA243 |  |
| pCA276.2 | pUV15-N-signal_sequence-mRFP-PBP-lipo_Mab_-strep | kan | CT94 |  |
| pCA308 | pUV15-dacB1-GFPmut3-strep | kan | CT94 |  |
| pCA324 | MOP-PBP-lipo_Mab_ (recode1)-strep | zeo | pCA243 |  |
| pCA325 | MOP-PBP-lipo_Mab_ (recode1)(S364A)-strep | zeo | pCA243 |  |
| pCA341 | piMyc-ftsZ-mNeonGreen | zeo | pCA243 |  |
| pCA342 | piMyc-dacB1-mNeonGreen | zeo | pCA243 |  |
| pCA344 | MOP-PbpB-strep | zeo | pCA243 |  |
| pCA353 | pUV15-N-signal_sequence-mRFP-PBP-lipo_Mab_(recode2)-strep | zeo | pCA264 |  |
| pCA354 | pUV15-N-signal_sequence-mRFP-PBP-lipo_Mab_-strep | zeo | pCA264 |  |
| pcr23 | pUV15-Tet-dCas9-cr23-*(rpoB)* | kan | CT296 |  |
| pcr25 | pUV15-Tet-dCas9-cr25-*(PBP-lipo_Mab_)* | kan | CT296 | 1 sgRNA strain used in the paper |
| pcr26 | pUV15-Tet-dCas9-cr26-*(PBP-lipo_Mab_)* | kan | CT296 |  |
| pcr36 | pUV15-Tet-dCas9-cr36-*(glnA2_Mab_)* | kan | CT296 |  |
| pcr40 | pUV15-Tet-dCas9-cr40-*(MAB_4471)* | kan | CT296 |  |
| pcr41 | pUV15-Tet-dCas9-cr41-*(gyrB_Mab_)* | kan | CT296 |  |
| pcr43 | pUV15-Tet-dCas9-cr43-*(secY_Mab_)* | kan | CT296 |  |
| pcr56 | pUV15-Tet-dCas9-cr56-*(PBP-lipo_Mab_)* | kan | CT296 |  |
| pcr61 | pUV15-Tet-dCas9-cr61-*(icd2_Mab_)* | kan | CT296 |  |
| pcr63 | pUV15-Tet-dCas9-cr63-*(sdhA_Mab_)* | kan | CT296 |  |
| pcr67 | pUV15-Tet-dCas9-cr67-*(ponA1_Mab_)* | kan | CT296 |  |
| pcr68 | pUV15-Tet-dCas9-cr68 *(ponA2_Mab_)* | kan | CT296 |  |
| pcr69 | pUV15-Tet-dCas9-cr69-*(pbpA_Mab_)* | kan | CT296 |  |
| pcr70 | pUV15-Tet-dCas9-cr70-*(pbpB_Mab_)* | kan | CT296 |  |
| pcr71 | pUV15-Tet-dCas9-cr71-*(dacB1_Mab_)* | kan | CT296 |  |
| pcr72 | pUV15-Tet-dCas9-cr72-*(dacB2_Mab_)* | kan | CT296 |  |
| pcr73 | pUV15-Tet-dCas9-cr73-*(carboxypeptidase_Mab_)* | kan | CT296 |  |
| pcr74 | pUV15-Tet-dCas9-cr74-*(VanY-type carboxypeptidase_Mab_)* | kan | CT296 |  |
| pcr86 | pUV15-Tet-dCas9-cr86-*(ponA1_Msm_)* | kan | CT295 |  |
| pcr87 | pUV15-Tet-dCas9-cr87-*(ponA1_Msm_)* | kan | CT295 |  |
| pcr88 | pUV15-Tet-dCas9-cr88-*(ponA2_Msm_)* | kan | CT295 |  |
| pcr89 | pUV15-Tet-dCas9-cr89-*(MSMEG_4384)* | kan | CT295 |  |
| pcr90 | pUV15-Tet-dCas9-cr90-*(pbpA_Msm_)* | kan | CT295 |  |
| pcr91 | pUV15-Tet-dCas9-cr91-*(pbpB_Msm_)* | kan | CT295 |  |
| pcr92 | pUV15-Tet-dCas9-cr92-*(PBP-lipo_Msm_)* | kan | CT295 |  |
| pcr93 | pUV15-Tet-dCas9-cr93-*(MSMEG_6319_Msm_)* | kan | CT295 |  |
| pcr94 | pUV15-Tet-dCas9-cr94-*(dacB1_Msm_)* | kan | CT295 |  |
| pcr95 | pUV15-Tet-dCas9-cr95-*(dacB2_Msm_)* | kan | CT295 |  |
| pcr96 | pUV15-Tet-dCas9-cr96-*(MSMEG_2432)* | kan | CT295 |  |
| pcr97 | pUV15-Tet-dCas9-cr97-*(carboxypeptidase_Msm_)* | kan | CT295 |  |
| pcr98 | pUV15-Tet-dCas9-cr98-*(VanY-carboxypeptidase_Msm_)* | kan | CT295 |  |
| pCRC1 | pUV15-Tet-dCas9-cr25+cr26 *(PBP-lipo)* | kan | cr25 | 2 sgRNA strain used in the paper |
| pCRC2 | pUV15-Tet-dCas9-cr25+cr26+cr56 *(PBP-lipo)* | kan | cr25 | 3 sgRNA strain used in the paper |
| pCRC5 | pUV15-Tet-dCas9-cr67-*(ponA1_Mab_)* + cr25 *(PBP-lipo_Mab_)* | kan | cr25 |  |
| pCRC6 | pUV15-Tet-dCas9-cr68 *(ponA2_Mab_)* + cr25 *(PBP-lipo_Mab_)* | kan | cr25 |  |
| pCRC7 | pUV15-Tet-dCas9-cr69-*(pbpA_Mab_)* + cr25 *(PBP-lipo_Mab_)* | kan | cr25 |  |
| pCRC8 | pUV15-Tet-dCas9-cr70-*(pbpB_Mab_)* + cr25 *(PBP-lipo_Mab_)* | kan | cr25 |  |
| pCRC9 | pUV15-Tet-dCas9-cr71-*(dacB1_Mab_)* + cr25 *(PBP-lipo_Mab_)* | kan | cr25 |  |
| pCRC10 | pUV15-Tet-dCas9-cr72-*(dacB2_Mab_)* + cr25 *(PBP-lipo_Mab_* | kan | cr25 |  |
| pCRC11 | pUV15-Tet-dCas9-cr73-*(carboxypeptidase_Mab_)* + cr25 *(PBP-lipo_Mab_)* | kan | cr25 |  |
| pCRC12 | pUV15-Tet-dCas9-cr74-*(VanY-type carboxypeptidase_Mab_)* + cr25 *(PBP-lipo_Mab_)* | kan | cr25 |  |
| pCRC13 | pUV15-Tet-dCas9-cr25 *(PBP-lipo_Mab_)* + empty guide | kan | cr25 |  |
