## Supplemental Table 9 for "High-density transposon mutagenesis in *Mycobacterium abscessus* identifies an essential penicillin-binding lipo-protein (PBP-lipo) involved in septal peptidoglycan synthesis and antibiotic sensitivity"

**Supplemental Table 9: Primers Used**

| **Primer Number** | **Sequence** | **Description** | **Purpose** |
| --- | --- | --- | --- |
| 938 | tccagcTTAGATATCaaggaggTATACATatgttcagacgtggatgggtcc | MAB_3167c (rc1) + pCA239 NdeI o/h F1 | pCA272 |
| 1050 | CGACATCGATAAGCTTGATTGCAGGATATCCTCGAGTCACTTCTCGAACTGGGGGTGGCTCCAGTCGGCCAAGTAGTCGGGCGGCAGC | MAB_3167c-strep + pCA243 XhoI o/h | “ “ |
| 693 | GCATGCTTAATTAAGAAGGAGATATACATatgttcagacgtggatgggtcc | MAB_3167c + pCT94 NdeI | pCA276.2/pCA341 |
| 1060 | GGCCATACCGCTGCCCGAGCCACCGCGCGGGGTACACGCGACAGC | ss-peptide(+3aa-linker2)-MAB_3167c-strep + mRFP o/h | “ “ |
| 1061 | GGTGGCTCGGGCAGCGGTATGGCCTCCTCCGAGGACGTC | mRFP o/h + linker2 | “ “ |
| 1105 | GGCGCGGGGTACCGCTGCCCGAGCCACCGCGGCCCTCGGCGCGCTCGTAC | linker2-mRFP + linker2-Mab_3167c-strep o/h | “ “ |
| 1059 | GGTGGCTCGGGCAGCGGTACCCCGCGCCCCGATGGGCCG | MAB_3167c-strep+ linker2 o/h | “ “ |
| 694 | TCTAGGGTCCCCAATTAATTAGCTAAAGCTTtcacttctcgaactgggggtggctccagtcggccaagtagtcgggcggcagc | MAB_3167c + pCT94 HindIII | “ “ |
| 893 | CATGCTTAATTAAGAAGGAGATATACATatgaccacctggactcctgcgc | MAB_3681 (dacB1) + CT302 NdeI o/h | pCA308 |
| 1142 | ACATCGAGGTCCCCCCCGAGCCGCCAGCCTGTCGGCGGTTGAGTGAGC | dacB1 + GFPmut3-linker o/h | “ “ |
| 1143 | GGCGGCTCGGGGGGGACCTCGATGTCGAAGGG | GFPmut3 + 3aa-linker | “ “ |
| 1144 | TCTAGGGTCCCCAATTAATTAGCTAAAGCTTTCACTTCTCGAACTGGGGGTGGCTCCAGTCGGTGATCCCCGCGGCGTTCACG | GFPmut3 + strep-tag + CT94 HindIII o/h | “ “ |
| 1159 | atccagcTTAGATATCaaggaggTATACATatgttcagacgtggatgggtcctgg | MAB_3167c + pCA243 o/h | pCA323 |
| 1160 | ttGAAGCTGGTGGCTTGCAAGCCTgaAaaCgcttggtccaggcccactttcgcg | MAB_3167c + PAM1 o/h | “ “ |
| 1161 | gcGttTtcAGGCTTGCAAGCCACCAGCTTCaaggccgcggtgaatggctcgca | MAB_3167c + PAM1 o/h | “ “ |
| 1162 | CGCAAACCACGCGTGGCTACCCCCAggGaactccgcttcgcccgtttttccgaag | MAB_3167c + PAM2 o/h | “ “ |
| 1163 | ttCccTGGGGGTAGCCACGCGTGGTTTGCGggataccgcggcgacatggccttcg | MAB_3167c + PAM2 o/h | “ “ |
| 1164 | tcgacatcgataagcttgattgcaggatatcCTCGAGtcacttctcgaactgggggtggctccagtcggccaagtagtcgggcggcagcg | MAB_3167c + pCA243 o/h | “ “ |
| 1165 | GGCACCGGCGGTGATGATCTTGAATGTGGCCCCAGGCGGATACAACCCGGTGG | MAB_3167c + S364A mutant o/h | pCA325 |
| 1166 | GCCACCACCGGGTTGTATCCGCCTGGGGCCACATTCAAGATCATCACCGCCGG | MAB_3167c + S364A mutant o/h | “ “ |
| cr.23a | GGGAGAACGACAGCGACATCGAGCC | sgRNA Fwd *rpoB_Mab_* | cr23 |
| cr.23b | AAACGGCTCGATGTCGCTGTCGTTC | sgRNA Rev *rpoB_Mab_* |  |
| cr.25a | GGGAGAACGACGTCGCCTGTAGTCC | sgRNA Fwd *PBP-lipo_Mab_* | cr25 |
| cr.25b | AAACGGACTACAGGCGACGTCGTTC | sgRNA Rev *PBP-lipo_Mab_* | “ “ |
| cr.26a | GGGAGCGAACCAGGCATGTGATCCGC | sgRNA Fwd *PBP-lipo_Mab_* | cr26 |
| cr.26b | AAACGCGGATCACATGCCTGGTTCGC | sgRNA Rev *PBP-lipo_Mab_* | “ “ |
| cr.36a | GGGAACTCGGCCCGCTTGTTGCGCAG | sgRNA Fwd  *glnA2_Mab_* | cr36 |
| cr.36b | AAACCTGCGCAACAAGCGGGCCGAGT | sgRNA Rev  *glnA2_Mab_* | “ “ |
| cr.40a | GGGAGGTTGCCGTAATCGGGCCGGT | sgRNA Fwd  *MAB_4471* | cr40 |
| cr.40b | AAACACCGGCCCGATTACGGCAACC | sgRNA Rev  *MAB_4471* | “ “ |
| cr.41a | GGGAGCTTGTACAACGGTGGCTGCGC | sgRNA Fwd  *gyrB_Mab_* | cr41 |
| cr.41b | AAACGCGCAGCCACCGTTGTACAAGC | sgRNA Rev  *gyrB_Mab_* | “ “ |
| cr.43a | GGGAGGGGTTGAAGGTGATCGACAC | sgRNA Fwd  *secY_Mab_* | cr43 |
| cr.43b | AAACGTGTCGATCACCTTCAACCCC | sgRNA Rev  *secY_Mab_* | “ “ |
| cr.56a | GGGAGCTTAGCGGGGTCTGTGCGCAGTG | sgRNA Fwd  *PBP-lipo_Mab_* | cr56 |
| cr.56b | AAACCACTGCGCACAGACCCCGCTAAGC | sgRNA Rev  *PBP-lipo_Mab_* | “ “ |
| cr.61a | GGGAGGCGAATGCCCGGATGATCGG | sgRNA Fwd  *icd2_Mab_* | cr61 |
| cr.61b | AAACCCGATCATCCGGGCATTCGCC | sgRNA Rev  *icd2_Mab_* | “ “ |
| cr.63a | GGGAGGTGGAATTCCATGTCCTCCA | sgRNA Fwd  *sdhA_Mab_* | cr63 |
| cr.63b | AAACTGGAGGACATGGAATTCCACC | sgRNA Rev  *sdhA_Mab_* | “ “ |
| cr.67a | GGGAGAACGAGAAGCCCGGGTTGCC | sgRNA Fwd  *ponA1_Mab_* | cr67 |
| cr.67b | AAACGGCAACCCGGGCTTCTCGTTC | sgRNA Rev  *ponA1_Mab_* | “ “ |
| cr.68a | GGGAGCCAGGTAGTCGAGCACGTAATC | sgRNA Fwd  *ponA2_Mab_* | cr68 |
| cr.68b | AAACGATTACGTGCTCGACTACCTGGC | sgRNA Rev  *ponA2_Mab_* | “ “ |
| cr.69a | GGGAGCCACGTGGGTCCCGGCCGGT | sgRNA Fwd  *pbpA_Mab_* | cr69 |
| cr.69b | AAACACCGGCCGGGACCCACGTGGC | sgRNA Rev  *pbpA_Mab_* | “ “ |
| cr.70a | GGGAACTGTCCAGTGAATCCTCAAGCC | sgRNA Fwd  *pbpB_Mab_* | cr70 |
| cr.70b | AAACGGCTTGAGGATTCACTGGACAGT | sgRNA Rev  *pbpB_Mab_* | “ “ |
| cr.71a | GGGAGCGGCGTGGAGAACCCATAGT | sgRNA Fwd  *dacB1_Mab_* | cr71 |
| cr.71b | AAACACTATGGGTTCTCCACGCCGC | sgRNA Rev  *dacB1_Mab_* | “ “ |
| cr.72a | GGGAACACCGGATTGGCCAGCGCGGC | sgRNA Fwd  *dacB2_Mab_* | cr72 |
| cr.72b | AAACGCCGCGCTGGCCAATCCGGTGT | sgRNA Rev  *dacB2_Mab_* | “ “ |
| cr.73a | GGGAACTTGTCACGGCGTCCACCGTGCC | sgRNA Fwd  *carboxypeptidase_Mab_* | cr73 |
| cr.73b | AAACGGCACGGTGGACGCCGTGACAAGT | sgRNA Rev  *carboxypeptidase_Mab_* | “ “ |
| cr.74a | GGGAGGGAACGCCAGCCCGACGTGA | sgRNA Fwd  *VanY-type carboxypeptidase_Mab_* | cr74 |
| cr.74b | AAACTCACGTCGGGCTGGCGTTCCC | sgRNA Rev  *VanY-type carboxypeptidase_Mab_* | “ “ |
| cr.86a | GGGAGCACCATCACGGCGACGGCGATC | sgRNA Fwd  *ponA1_Msm_* | cr86 |
| cr.86b | AAACGATCGCCGTCGCCGTGATGGTGC | sgRNA Rev  *ponA1_Msm_* | “ “ |
| cr.87a | GGGAGCGTAGTCGGGGGGCACCGTCGG | sgRNA Fwd  *ponA1_Msm_* | cr87 |
| cr.87b | AAACCCGACGGTGCCCCCCGACTACGC | sgRNA Rev  *ponA1_Msm_* | “ “ |
| cr.88a | GGGAGCCAGGTACTCCAGCACGTAGTC | sgRNA Fwd  *ponA2_Msm_* | cr88 |
| cr.88b | AAACGACTACGTGCTGGAGTACCTGGC | sgRNA Rev  *ponA2_Msm_* | “ “ |
| cr.89a | GGGAGCTGGTGGACCGCACCAGGCCCG | sgRNA Fwd *MSMEG_4384* | cr89 |
| cr.89b | AAACCGGGCCTGGTGCGGTCCACCAGC | sgRNA Rev  *MSMEG_4384* | “ “ |
| cr.90a | GGGAGCCGCCGCGCGGGTCGCGGCCGGT | sgRNA Fwd  *pbpA_Msm_* | cr90 |
| cr.90b | AAACACCGGCCGCGACCCGCGCGGCGGC | sgRNA Rev  *pbpA_Msm_* | “ “ |
| cr.91a | GGGAGCCCGAACTTGCGCAGCATCTC | sgRNA Fwd  *pbpB_Msm_* | cr91 |
| cr.91b | AAACGAGATGCTGCGCAAGTTCGGGC | sgRNA Rev  *pbpB_Msm_* | “ “ |
| cr.92a | GGGAGTAGTGGTAGATCTGCCCCGGCA | sgRNA Fwd  *PBP-lipo_Msm_* | cr92 |
| cr.92b | AAACTGCCGGGGCAGATCTACCACTAC | sgRNA Rev  *PBP-lipo_Msm_* | “ “ |
| cr.93a | GGGAGCAATTCGCTGTCCTCGCTGTA | sgRNA Fwd  *MSMEG_6319_Msm_* | cr93 |
| cr.93b | AAACTACAGCGAGGACAGCGAATTGC | sgRNA Rev  *MSMEG_6319_Msm_* | “ “ |
| cr.94a | GGGAGCGCTGATTGAGTTGGCGCGCAC | sgRNA Fwd  *dacB1_Msm_* | cr94 |
| cr.94b | AAACGTGCGCGCCAACTCAATCAGCGC | sgRNA Rev  *dacB1_Msm_* | “ “ |
| cr.95a | GGGAAGACCGGGTCGGCCATGGCGGC | sgRNA Fwd  *dacB2_Msm_* | cr95 |
| cr.95b | AAACGCCGCCATGGCCGACCCGGTCT | sgRNA Rev  *dacB2_Msm_* | “ “ |
| cr.96a | GGGAATCACCGGATAGTTCAGCGCCGC | sgRNA Fwd  *MSMEG_2432* | cr96 |
| cr.96b | AAACGCGGCGCTGAACTATCCGGTGAT | sgRNA Rev  *MSMEG_2432* | “ “ |
| cr.97a | GGGAGTGGGCTGTGTGCGCCCACCGT | sgRNA Fwd  *carboxypeptidase_Msm_* | cr97 |
| cr.97b | AAACACGGTGGGCGCACACAGCCCAC | sgRNA Rev  *carboxypeptidase_Msm_* | “ “ |
| cr.98a | GGGAGCCGGTTGCTGCAGGTCGAACGG | sgRNA Fwd  *VanY-carboxypeptidase_Msm_* | cr98 |
| cr.98b | AAACCCGTTCGACCTGCAGCAACCGGC | sgRNA Rev  *VanY-carboxypeptidase_Msm_* | “ “ |
| 855 | AGACTCGCTCTTCCGGAtctgaccagggaaaatagc | 1Ab fragment - Golden Gate Cloning pCT296 (SapI) + NO Handle | Golden Gate Cloning 2sgRNA Fwd |
| 856 | AGACTCGCTCTTCCCTGAAAATAAAAAAGGGGACCTCTAG | 1Ab fragment - Golden Gate Cloning pCT296 (SapI) + NO Handle | Golden Gate Cloning 2sgRNA Rev |
| 857 | CTCCTTGCTCTTCCGGAtctgaccagggaaaatagc | 2A - Golden Gate Handle F1 pCT296 (SapI) + no handle | Golden Gate Clonin3 2sgRNA Fwd 1 |
| 858 | CTCCTTGCTCTTCCaAAAATAAAAAAGGGGACCTCTAG | 2A - Golden Gate Handle F1 pCT296 (SapI) + no handle | Golden Gate Clonin3 2sgRNA Rev 1 |
| 859 | CTCCTTGCTCTTCCTTtctgaccagggaaaatagc | 2B - Golden Gate Handle F1 pCT296 (SapI) + no handle | Golden Gate Clonin3 2sgRNA Fwd 2 |
| 860 | CTCCTTGCTCTTCCCTGAAAATAAAAAAGGGGACCTCTAG | 2B - Golden Gate Handle F1 pCT296 (SapI) + no handle | Golden Gate Clonin3 2sgRNA Rev 2 |
| 978 | ggtccggtggcggagaagtttt | PBP-lipo_Mab_ qPCR primers |  |
| 979 | ttcgttgaccgtcgcccgctta | " " |  |
| 980 | ttcggccctacgacgacaccat | " " |  |
| 981 | ggccacgggtattgacggcatt | " " |  |
| 982 | gggttgtatccgcctgggtcga | " " |  |
| 983 | gaacggcgtgaccagcaccttg | " " |  |
| 984 | CGGAATGATGCGCGAGACGGT | " " |  |
| 985 | GCGGCAGCGACTGGAACATCAC | " " |  |
| 986 | gccacgaaagcagctccggtaa | sigA_Mab_ qPCR primers |  |
| 987 | tctcgcccttctcggtcagctc | " " |  |
| 988 | atcgagcccggcgatgacttgg | " " |  |
| 989 | ttcgatgaagtcgccgagctgc | " " |  |
| 990 | gcatggcgttcctggacctgat | " " |  |
| 991 | agcacctgtgaacggctcgg | " " |  |
| 916 | agcgctcacaattcggatccagcTTAGATATCaaggaggTATACATatggcaactcgaacatcaccagtc | MSMEG_2584 + pJW460 EcoRV o/h | pCA242 |
| 919 | TCGATAAGCTTGATTGCAGGATATCCTCGAGTCACAGGTCTTCCTCGCTGATCAGCTTCTGCTCGACGAGGTAGTCCGCGGGC | MSMEG_2584--myc + pCA239/240 XhoI o/h | " " |
| 868 | CGCCGAGCGGGTCCGCGACGCC | 500bp u/s MSMEG_2584 | MSMEG_2584 recombineering construct |
| 869 | TACCCAACTTAATCGCCTTGCAGCTGAGACTGATGGTAAGAACGTTGC | zeo LF + 500bp u/s MSMEG_2584 o/h | " " |
| 870 | GCAACGTTCTTACCATCAGTCTCAGCTGCAAGGCGATTAAGTTGGGTA | 500bp u/s MSMEG_2584 + zeo LF o/h | " " |
| 976 | CCACATCGGCCGAGATCACACGCGAAAACAGCTATGACCATGATTACGCCA | zeoR + MSMEG_2584 o/h | " " |
| 977 | GGCGTAATCATGGTCATAGCTGTTTTCGCGTGTGATCTCGGCCGATG | 500bp d/s MSMEG_2584 + zeoR o/h | " " |
| 971 | GGGCCGCGGATCTCCACCGAGC | MSMEG_2584 600bp d/s s.primer | " " |
| 1024 | actagtgttCTGCATGCACTCTAGAAATATTCTCCAGCCCTGACCTGCCCACC | ftsZ natP + pCA243 o/h | pCA269 |
| 1025 | CCCCCTTCGACATCGAGGTCCCCCCCATGGTCGCCTTCCTCCCTGGTTGC | ftsZ natP + GFPmut3 o/h | " " |
| 1026 | GCAGCAACCAGGGAGGAAGGCGACCATGGGGGGGACCTCGATGTCG | GFPmut3 + natP o/h | " " |
| 1021 | CGAGCCGCCGGTGATCCCCGCGGCGTTCACG | GFPmut3 + linker o/h | " " |
| 1022 | CGCCGCGGGGATCACCGGCGGCTCGatgacgcctccacacaactacc | ftsZ + GFPmut3 o/h | " " |
| 1023 | TCTAGGGTCCCCAATTAATTAGCTAGATATCTCAtcgccgcatgaagggcggc | ftsZ + pCA243 o/h | " " |
| 1028 | GGAATTGCGCGAGTTCTTTGCG | LF-Rv2864c 550bp u/s | Rv2864c recombineering construct |
| 1029 | TACCCAACTTAATCGCCTTGCAGCCGCGGCTCGCCCCTCCTCGTTTG | LF-Rv2864c 550bp + zeoR o/h | " " |
| 1030 | CAAACGAGGAGGGGCGAGCCGCGGCTGCAAGGCGATTAAGTTGGGTA | zeoR + LF-Rv2864c 500bp u/s o/h | " " |
| 1031 | GTCACAGTTCTTAACATCAGCAACGAAACAGCTATGACCATGATTACGCCA | zeoR + LF-Rv2864c 500bp d/s o/h | " " |
| 1032 | GGCGTAATCATGGTCATAGCTGTTTCGTTGCTGATGTTAAGAACTGTG | LF-RV2864c 500bp d/s + zeoR o/h | " " |
| 1033 | GCACGGTGTAAGGCACCGCTCAG | LF-Rv2864c 550bp d/s | " " |
| 1191 | ATGCCTGGCAGTCGATCGTACGCTAGTTAACGTTTAAACGGATCGTCGCACCG | PMyc + pCA243 HpaI o/h | pCA341 |
| 1192 | cgaggtagttgtgtggaggcgtcatATGTATACCTCCTGATATCTAATTGC | PMyc + ftsZ o/h | " " |
| 1193 | GCAATTAGATATCAGGAGGTATACATatgacgcctccacacaactacc | ftsZ + pMyc o/h | " " |
| 1194 | TGTTATCCTCCTCGCCCTTGCTCACCGAGCCGCCtcgccgcatgaagggcggcacg | ftsZ + mNeonGreen o/h | " " |
| 1195 | cgacgtgccgcccttcatgcggcgaGGCGGCTCGGTGAGCAAGGGCGAGGAGGATAAC | mNeonGreen + ftsZ o/h | " " |
| 1196 | gatgcctggcagtcgatcgtacgctaCTCGAGgttaacTTACTTGTACAGCTCGTCCATGC | mNeonGreen + pCA243 HpaI o/h | " " |
| 1197 | CCGCAATTAGATATCAGGAGGTATACATatgaccacctggactcctgcgc | dacB1 + pCA341 NdeI o/h | pCA342 |
| 1198 | TGTTATCCTCCTCGCCCTTGCTCACCGAGCCGCCagcctgtcggcggttgagtgagc | dacB1 + mNeonGreen o/h | " " |
| 1199 | gcgctcactcaaccgccgacaggctGGCGGCTCGGTGAGCAAGGGCGAGGAGGATAAC | mNeonGreen + dacB1 o/h | " " |
| 1203 | ccagcTTAGATATCaaggaggTATACATgtgaaccggcaggatcggtccc | pbpB + pCA243 NdeI o/h | pCA344 |
| 1204 | tcgataagcttgattgcaggatatcCTCGAGtcacttctcgaactgggggtggctccagtcggtcgcctgcaggatcagtggc | pbpB + pCA243 XhoI o/h | “ " |
| 1058 | ggcgcggggtACCGCTGCCCGAGCCACCgcggccctcggcgcgctcgtac | linker2-mRFP + linker2-Mab_3167c-strep o/h | pCA353 |
| 1059 | GGTGGCTCGGGCAGCGGTaccccgcgccccgatgggccg | linker2-Mab_3167c-strep + linker2-mRFP + o/h | " " |
| 1066 | GAAGCTGGTGGCTTGCAAGCCTgaAaaCgcttggtccaggcccactttcg | PBP-lipo + recode scramble 1 | " " |
| 1067 | GttTtcAGGCTTGCAAGCCACCAGCTTCaaggccgcggtgaatggctcgc | PBP-lipo + recode scramble 1 | " " |
| 1153 | CCTCTAGGGTCCCCAATTAATTAGCTAGATATCtcacttctcgaactgggggtggctccagtcggccaagtagtcgggcggcagc | PBP-lipo-strep + pCA264 o/h | pCA353/  pCA354 |
| *Mab* qPCR Primers | | | |
| 1171 | TGAAGAAGATCCGCAGTGATG | pbpB Set 1 |  |
| 1172 | CTCGGGAAACTCCTTGGTTATC | pbpB Set 1 |  |
| 1173 | CTGACTTTCCAGCCGAAGAA | pbpB Set 2 |  |
| 1174 | CTTATCGGCGTCACTCTTCC | pbpB Set 2 |  |
| 1175 | CCTCGGCCTGTTCTACAAATAC | dacB1 Set 1 |  |
| 1176 | GTTGTCGTTCGTCATCTCATAGG | dacB1 Set 1 |  |
| 1177 | GGCACACCTTCTCGACTATG | dacB1 Set 2 |  |
| 1178 | GCATGCGGTTTGGTCAAC | dacB1 Set 2 |  |
| 1187 | CCGTTCGATACTTCCTCTCTTG | Putative VanY-type carboxypeptidase Set 1 |  |
| 1188 | CGACGTGATCAGAAGGGTTAC | Putative VanY-type carboxypeptidase Set 1 |  |
| 1189 | GGTTGGATCCTTCGCTACTG | Putative VanY-type carboxypeptidase Set 2 |  |
| 1190 | CCATACGTCTGTACCGCATC | Putative VanY-type carboxypeptidase Set 2 |  |
| *Msm* qPCR Primers | | | |
| 1205 | CAGATCCTGTCGCTCCGAACCGACT | pbpB (Set 1) MSMEG_4233 |  |
| 1206 | GAACAACACGCCGGACGCCAAGACC | pbpB (Set 1) MSMEG_4233 |  |
| 1207 | TGTTGCGATCGATGATGCTGCCGC | pbpB (Set 2) MSMEG_4233 |  |
| 1208 | TCGTCTTTCGTGTTCCGGCACCG | pbpB (Set 2) MSMEG_4233 |  |
| 1209 | CTGGGTGTGGACGATGTCGGCGAAC | dacB1 (Set 1) MSMEG_1661 |  |
| 1210 | AGCTCAACACGCTGGCCGCCAA | dacB1 (Set 1) MSMEG_1661 |  |
| 1211 | GCTGAACGAACTCAACCTCAA | dacB1 (Set 2) MSMEG_1661 |  |
| 1212 | GCAGATCGTTGATCGTGTACTG | dacB1 (Set 2) MSMEG_1661 |  |
| 1213 | GCGAGTACGTGCAGACC | VanY carboxypeptidase (Set 1) MSMEG_1900 |  |
| 1214 | CAGGATTCGTTGGCGTAGAT | VanY carboxypeptidase (Set 1) MSMEG_1900 |  |
| 1215 | ACAACGCGGTTCAGACCTA | VanY carboxypeptidase (Set 2) MSMEG_1900 |  |
| 1216 | CGATGTCGACGGCCTCA | VanY carboxypeptidase (Set 2) MSMEG_1900 |  |
| 1217 | CCGGTTCGACGTTCAAGAT | PBP-lipo (Set 1) MSMEG_2584 |  |
| 1218 | GATCGAAACCGCCGTAGTT | PBP-lipo (Set 1) MSMEG_2584 |  |
| 1219 | GAGCCTCATCACGCTCAAG | PBP-lipo (Set 2) MSMEG_2584 |  |
| 1220 | TCGGTGGGCAGCAATTC | PBP-lipo (Set 2) MSMEG_2584 |  |

o/h = overhang (used in Gibson cloning)
